# Nanobodies against the S2 region of the spike protein potently neutralize SARS-CoV-2 viruses and show resistance to virus escape

**DOI:** 10.64898/2026.01.08.698330

**Authors:** John Clarke, Luke Jones, Imogen Buckle, Parul Sharma, Emily Park, Anja Kipar, Adam Kirby, Daniele Mega, Siva Ramadurai, Aparajita Karmakar, Katy Cornish, Leah McCaffrey, Hannah Campaigne, Lauren E.-A. Eyssen, William S. James, James P. Stewart, Miles W. Carroll, Raymond J. Owens

## Abstract

Entry of coronaviruses into cells is mediated by the viral spike (S) glycoproteins each consisting of S1 receptor binding and S2 membrane fusion subunits. The sequence of the S2 region is very highly conserved amongst variants of SARS-CoV-2 and compared to the S1 unit shares significant sequence identity amongst different beta-coronavirus lineages. By targeting the S2 of SARS-CoV-2 we have identified two selective and potent neutralizing nanobodies (BA.1-C2 and BA.1-D3) that bind to two different quaternary epitopes in the S2 formed by the Heptad Repeat 2 (HR2) trimer at the base of the spike protein. The HR2 sequence is identical in SARS-CoV and SARS-CoV-2 but differs in other beta-coronaviruses explaining the lack of binding to the spike proteins of MERS-CoV or HuCoV-OC43. No viral escape was observed following serial passaging of SARS-CoV-2 (JN.1) with a combination of BA.1-C2 and BA.1-D3 and the most potent of these nanobodies reduced viral load in the hamster model of COVID-19, following intranasal administration. Overall, the results show the value of nanobody technology for identifying novel neutralising epitopes in the S2 region of beta-coronaviruses with potential for the development of new selective anti-viral agents.

## Introduction

The spike [1, 2] proteins of SARS-CoV-2 and other beta-coronaviruses form trimers on the surface of the virions and comprise two subunits, S1 and S2. Following receptor binding through the S1 region, the spike proteins undergo cleavage between S1 and S2, followed by conformational re-arrangement of the S2 subunit to expose the fusion peptide. This leads to fusion of the host and viral membranes and internalisation of the viral genome [3]. The amino acid sequences of the S2 region of SARS-CoV-2 variants of concern are highly conserved (98-100 % identity) and show significantly greater sequence similarity to other beta-coronaviruses compared to the S1 region (46% vs 29%). Therefore, targeting the S2 region for the development of vaccines and antibody-based therapeutics may lead to greater resistance to escape mutants and have the prospect of generating broadly cross-reactive antibodies against current and possible future beta-coronaviruses.

Several neutralising monoclonal antibodies have been isolated from either COVID-19 patients [4, 5] or mice immunised with spike proteins [6, 7] that bind to a linear epitope in the S2 domain of the spike protein around the stem-helix region encompassing residues 1142 - 1247. This sequence is highly conserved between beta-coronaviruses and some of these anti-S2 antibodies showed broad neutralisation activity against SARS-CoV-2, MERS-CoV and HuCoV-OC43 [6, 7]. A second class of human monoclonal antibodies that bind to the fusion peptide have also been reported which showed neutralisation activity [8, 9]. The sequence that includes the fusion peptide has been shown to be highly immunogenic from a survey of patient sera using a tiled peptide array covering the spike protein sequence [10].

Single domain antibodies (nanobodies), the vast majority of which bind to the Receptor Binding Domain (RBD) of S1, have proved effective in neutralising SARS-CoV-2 viruses both *in vitro* and in animal models of COVID-19 [11, 12]. By contrast, there are only a few reports of the isolation of nanobodies that bind to the S2 region of the spike protein [13–17]. In some cases, these have identified neutralising epitopes that have been localised to either the stem-helix (residues 1130 – 1149) [14] also identified by human monoclonal antibodies from convalescent patients [4, 5], or the Heptad Repeat-2 (HR2 residues 1163-1202) of the spike protein [15, 18]. These regions of the S2 are part of the fusion machinery of the spike protein and undergo major conformational rearrangement in the transition from pre-to post-fusion states [19]. The HR2 sequence is conserved amongst SARS-CoV-2 Omicron variants but diverges in other beta-coronaviruses (e.g. HuCoV-OC43 and MERS-CoV).

We have produced a nanobody phage display library from a llama immunized with the stabilized pre-fusion trimers of a mixture of SARS-CoV-2 Omicron BA.1, MERS-CoV and HuCoV-OC43. By screening the library with truncated spike proteins of SARS-CoV-2 we have identified nanobodies that selectively neutralise key Omicron variants of the virus and in combination mitigate the effects of virus escape.

## Results

### Screening for VHHs to the S2 spike region of SARS-CoV-2

A VHH phage display library was constructed from the peripheral blood mononuclear cells of a llama immunised with a cocktail of three beta-coronavirus spike proteins (SARS-CoV-2 Omicron BA.1, MERS-CoV and HCoV-OC43). The spike proteins were produced as stabilised trimers by substitution of proline residues into the S2 sequences and fusion to the T4 fibritin foldon sequence at the carboxy terminus [20–22]. The primary phage display library was screened with the trimerized and proline stabilised S2 fragment of the SARS-CoV-2 Omicron BA.1 spike protein (residues 681 to 1200) from which VHH binders were identified by phage ELISA using biotinylated BA.1 S2 to coat plates. The sub-library generated by panning with BA.1 S2 was re-screened with the corresponding S2 fragment of MERS-CoV (residues 749 to 1291), and further binders identified by MERS S2 phage ELISA (Fig. 1a). A second display sub-library was generated by panning the primary library with the spike protein of HuCoV-OC43 and this in turn was panned with MERS S2. Sequencing and clustering of phage ELISA positive clones from the three panning experiments gave a total of 24 different VHH sequences, 13 of which were unique to the BA.1 S2 screen, 11 were found in the two sub-libraries screened with MERS S2, of which 5 were common to both screening with BA.1 S2 and MERS S2 (Fig. 1b Supplementary Table 1).

**Figure 1.**
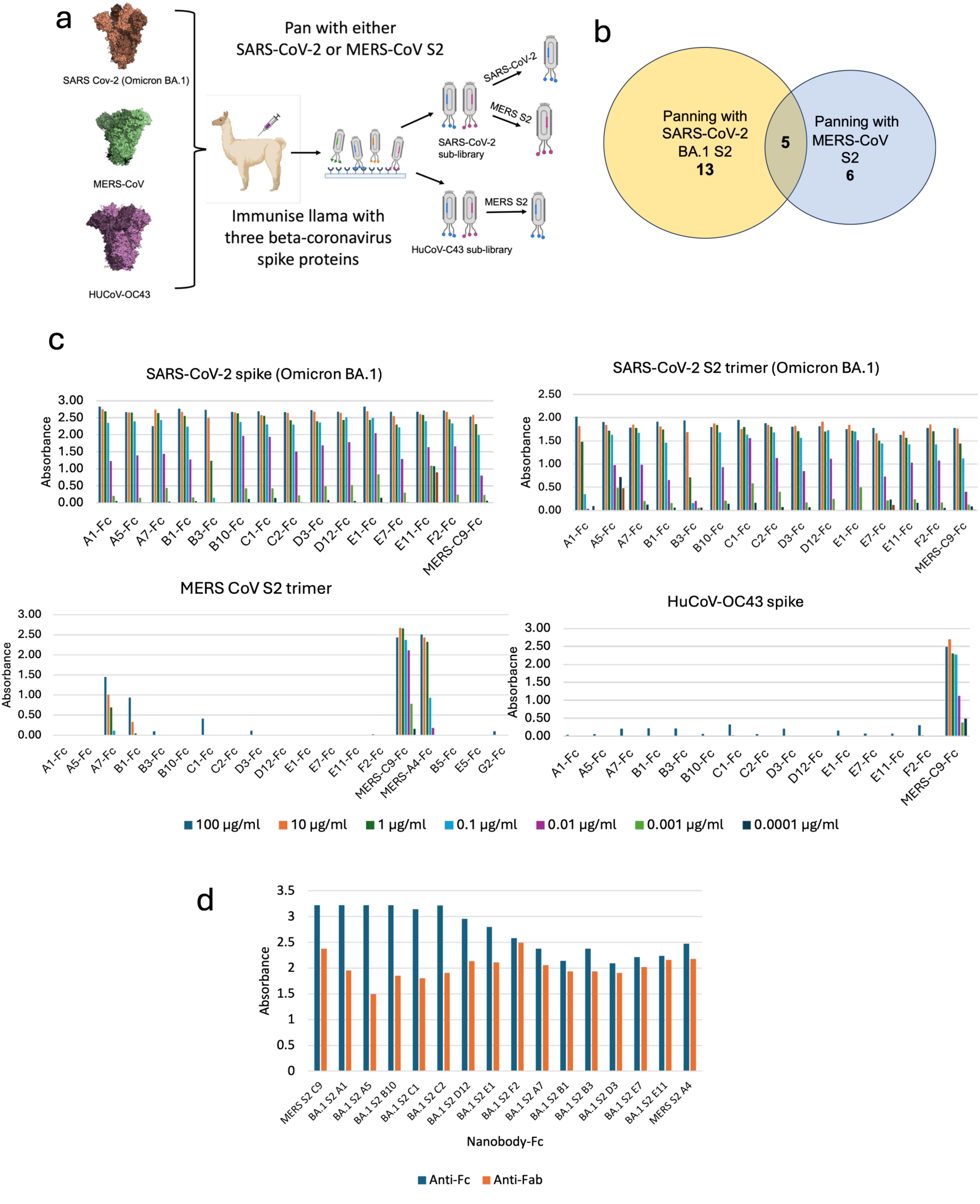
Nanobodies to the S2 region of beta-coronaviruses. **a** Schematic diagram summarising the immunization and panning of phage display libraries. **b** Venn diagram showing the number of nanobody sequences that were either common or unique from panning the spike library with either SARS-CoV-2 (Omicron BA.1) or MERS-CoV S2 trimers. **c** ELISA screening of nanobody-Fc fusions for binding to spike and S2 proteins detected by an HRP-anti-HuIgG1Fc reporter. **d** Binding of Nanobody-Fcs to SARS-CoV-2 BA.1 S2 trimer in the presence of the Fab fragment of anti-S2 monoclonal antibody, B6, [7] with Nanobody-Fcs detected with an anti-human IgG1-HRP antibody conjugate and B6 with an anti-mouse light chain HRP conjugate.

### Binding and neutralization profiles of anti-S2 nanobodies

The most abundant VHHs, including all those that were common to both panning with SARS-CoV-2 S2 and MERS-CoV S2 were selected for the construction of human IgG1 Fc fusions, expressed in Expi293F cells and purified by a combination of IMAC-SEC. The binding of the Fc fusions to either the spike and/or S2 proteins of SARS-CoV-2 (Omicron BA.1) MERS-CoV and HuCoV-OC43 was assayed by ELISA. The results showed that all nanobodies expressed as human IgG1 Fc fusions bound to the S2 subunit of both SARS-CoV-2 (BA.1) presumptively to common conserved epitopes (Fig. 1b). Only four of these nanobody Fc fusions (Nb-Fc) also bound to MERS-CoV S2, (MERS-C9, MERS-A4, BA.1-A7, BA.1-B1), of which only two showed binding to the spike protein of HuCoV-OC43 (MERS-C9 and MERS-A4). To assess whether any of the anti-S2 nanobodies bound to the same S2 epitopes previously identified for neutralising monoclonal antibodies, two reference antibodies that have been reported to bind to the stem-helix; B6, [7] and fusion peptide, CoV-91.27 [8] respectively, were cloned and expressed as Fab fragments. No binding of CoV-91.27 to SARS-CoV-2 S2 was observed, presumably due to a lack of exposure of the fusion peptide in the stabilised S2 trimer and confirming that it corresponds to a pre-fusion state. Given that the nanobodies bound to the S2 trimer we deduce that none bind to the fusion peptide. All the nanobodies bound to BA.1 S2 trimer in the presence of the stem-helix binding Fab B6, indicating that they do not bind to the stem-helix sequence (Fig.1c).

The nanobody Fc fusions were screened for neutralisation of SARS-CoV-2 virus (Prototypical strain Vic01 and Omicron BA.1) in authentic virus micro-neutralisation assays. Four nanobodies showed half maximal neutralisation titres (NT_50_) of less than 100 nM, with BA.1-C2 and D3 the most potent, both with NT50s of 1.6 nM against BA.1 (Table 1). These results showed that not all the epitopes recognised by the anti-S2 VHHs are neutralising, presumably reflecting the location of the binding sites on the S2 subunit.

**Table 1.** Screening nanobody Fc fusions for virus neutralisation activity.

| Nanobody-Fc | SARS-CoV-2<br>Vic01<br>NT<sub>50</sub> (nM) | SARS-CoV-2<br>Omicron BA.1<br>NT<sub>50</sub> (nM) |
| --- | --- | --- |
| BA.1 A1-Fc | >1000 | >1000 |
| BA.1 A5-Fc | >1000 | >500 |
| BA.1 A7-Fc | NI | NI |
| BA.1 B1-Fc | >1000 | >1000 |
| BA.1 B3-Fc | >1000 | >1000 |
| BA.1 B10-Fc | 60 | 60 |
| BA.1 C1-Fc | NI | >1000 |
| BA.1 C2-Fc | 0.7 | 1.6 |
| BA.1 D3-Fc | 20 | 1.6 |
| BA.1 D12-Fc | 40 | 20 |
| BA.1 E1-Fc | NI | NI |
| BA.1 E7-Fc | >1000 | >1000 |
| BA.1 E11-Fc | >1000 | >1000 |
| BA.1 F2-Fc | NI | NI |
| MERS C9-Fc | >1000 | >1000 |
| MERS A4-Fc | >1000 | >1000 |

NI = no inhibition > 10,000 nM
ND = not determined

The binding affinities of the four most potent neutralising anti-SARS-CoV-2 nanobodies, BA.1-B10, BA.1-C2, BA.1-D3, and BA.1-D12, were measured by BLI. The results confirmed tight binding of BA.1-B10, BA.1-D3, BA.1-D12, BA.1-C2 to SARS-CoV-2 (Fig. 2a). There was no direct correlation between apparent binding affinity and neutralization amongst these four nanobodies, with the most potent neutralizing nanobody, BA.1-C2 showing lowest affinity. A competition experiment was then carried out by BLI to determine whether the four nanobodies bound to the same or different epitopes in the S2 region of the SARS-CoV-2 spike protein. The results showed that BA.1-C2 and BA.1-D12 bound to the same or overlapping epitopes whereas BA.1-D3 and BA.1-B10 each bound to a different epitope (Fig.2b).

**Figure 2.**
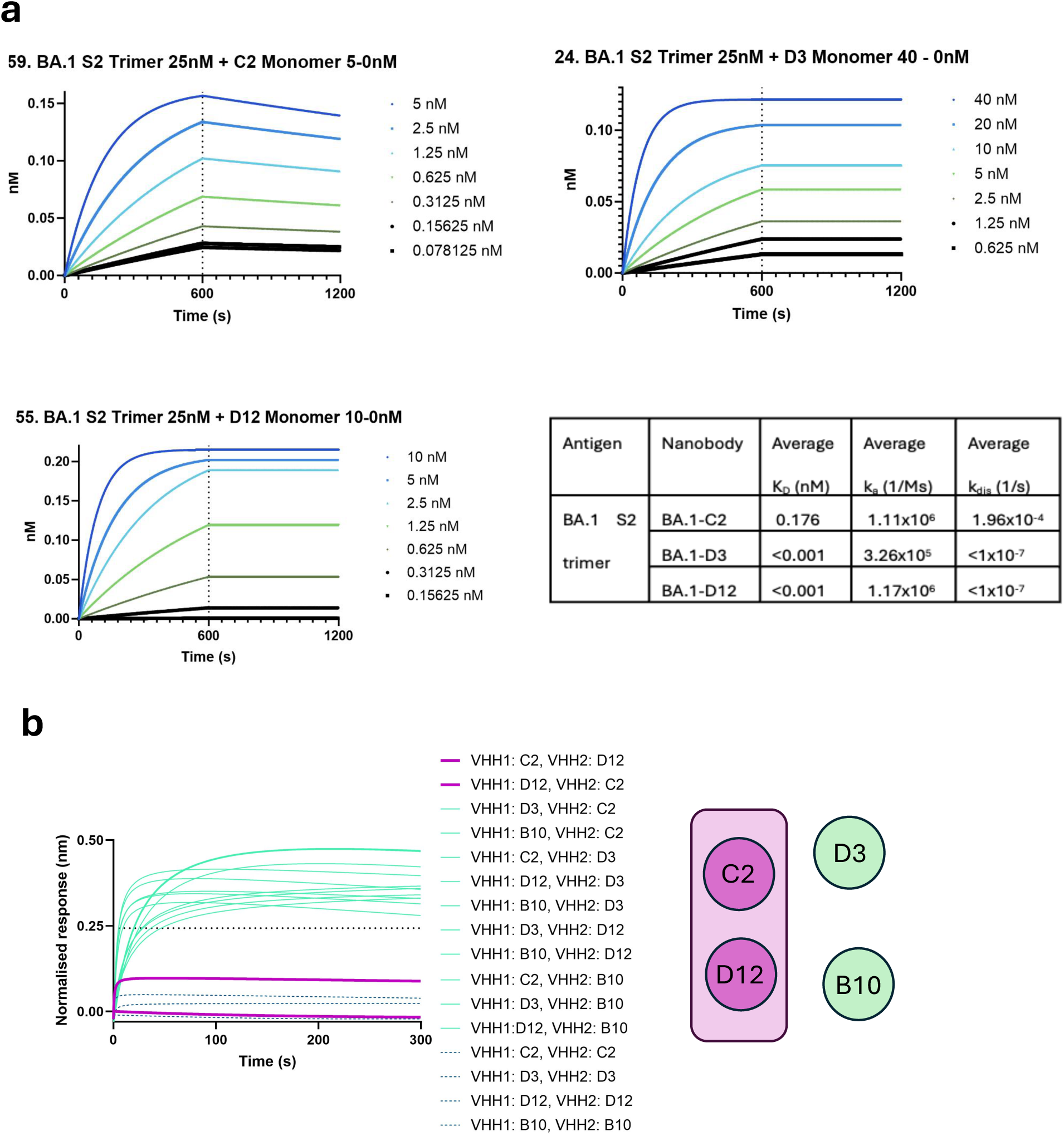
Kinetic binding affinities of monomeric nanobodies. **a** BLI sensorgrams and table of calculated affinities of the binding kinetics of BA.1-C2, BA.1-D3, BA.1-D12 and BA.1-B10 to SARS-CoV-2 Omicron BA.1 S2 trimer (25 mM) biotinylated and immobilized onto streptavidin sensors. The results are representative of two independent assays. **b** Epitope binning results for nanobody monomers binding to SARS-CoV-2 BA.1 S2 trimers measured by Biolayer Interferometry (BLI). The compiled BLI sensorgrams are coloured according to whether pairwise combinations of nanobodies are competitive (purple, bold), non-competitive (green) and self-v-self (blue, dashes). The horizontal black dotted line is the threshold set for determining competitive/non-competitive, set at 50% of the Min-Max range (= 0.24315). **c** Schematic representation of the results of the competition BLI.

### Neutralisation of SARS-CoV-2 Omicron variants by BA.1-C2 and BA.1-D3

For single domain antibodies that target the RBD of beta-coronaviruses dimeric and trimeric versions have been shown to have increased neutralisation activity compared to the corresponding monomers [23–25]. Previously, trimeric version of nanobodies that bound to the Receptor Binding Domain (RBD) of the spike protein of SARS-CoV-2 were been shown to potently neutralise Omicron viral variants both *in vitro* and in an animal of COVID-19 [26]. Therefore, trimeric versions of the two most potent neutralising nanobodies, BA.1-C2 and BA.1-D3, were constructed by joining VHHs end-to-end with six residue linkers comprising alternating glycine and serine residues, [GS]_3._ Neutralisation activity was compared to the corresponding monomers against four Omicron variants of SARS-CoV-2 (BA.1, EG.1, JN.1 and XBB1.5). The results showed that the monomers and trimers were equipotent against all Omicron variants and that in contrast to nanobodies targeting the RBD, there was no gain in activity by the trimeric versions (Fig. 3). The BA.1-C2 nanobody was approximately 10-fold more potent than the BA.1-D3 in both monomer and trimer formats and unexpectedly showed greater activity than the Fc fusion version previously tested (Table 1). The results indicate that only one subunit of the trimer is required for activity and that linking to an IgG Fc in some way negatively affects the interaction with the virus. Nanobody binding presumably disrupts the conformational transition of the spike protein from a pre-fusion trimer to the post-fusion state and hence prevents virus entry into the cells. The neutralisation results of the SARS-CoV-2 Omicron sub-variants, including the recent JN.1 strain which is resistant to treatment with many monoclonal antibodies [27], showed that targeting the highly conserved S2 region of the spike protein leads to broad reactivity against the Omicron variants of the virus.

**Figure 3.**
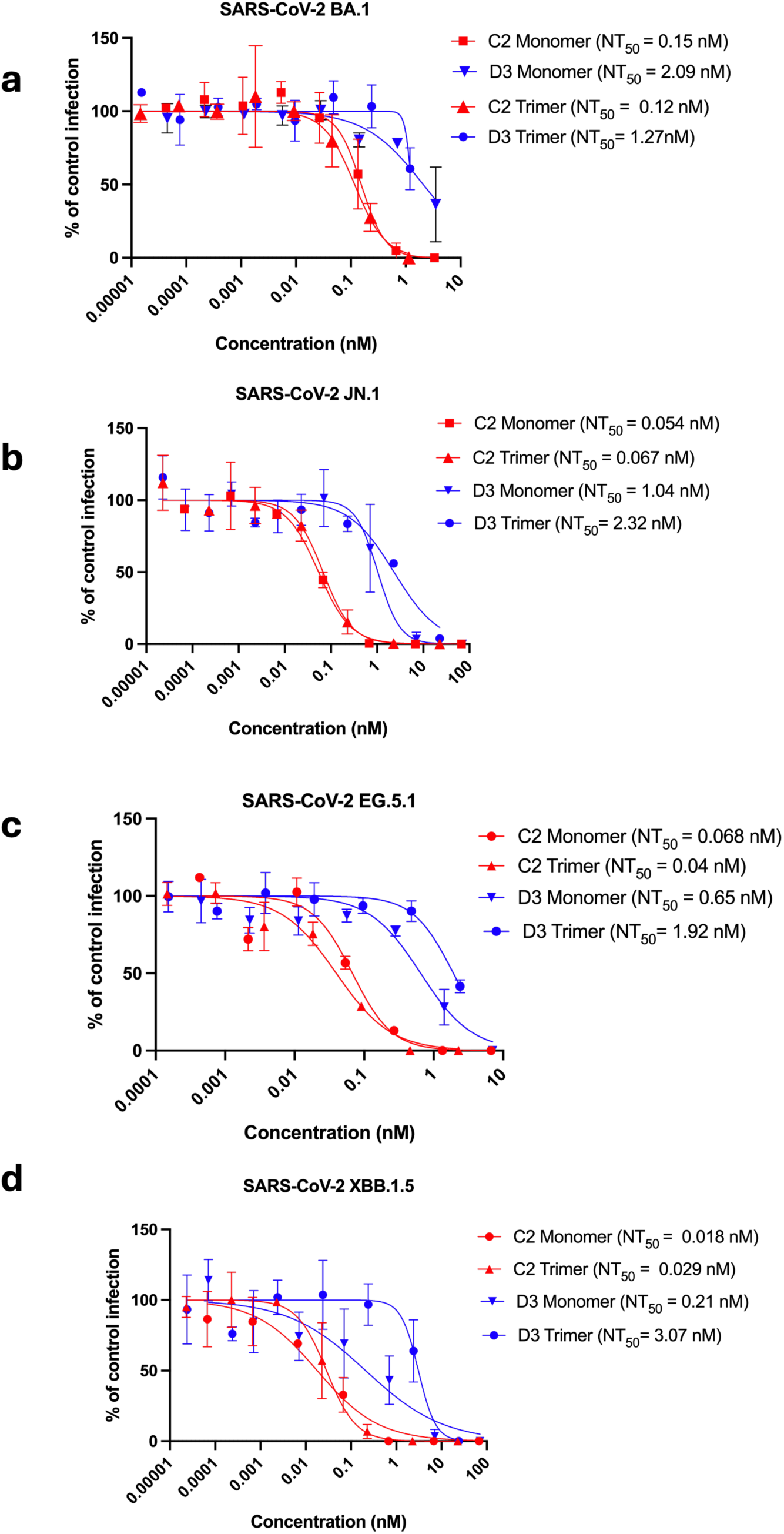
Neutralisation of SARS-CoV-2 Omicron variants. Neutralisation curve of SARS-CoV-2 Omicron variants measured in MNA **a** BA.1, **b** JN.1, **c** EG5.1 and **d** XBB.1.5. Data are shown as mean (n =2) +/- 95% Cl.

### Analysis of virus escape by BA.1-C2 and BA.1-D3 nanobodies

A viral escape study was carried out as a first approach to determining where the SARS-CoV-2 neutralising nanobodies, BA.1-C2 and D3 bound to the spike protein. VeroE6 cells, expressing TMPRSS2, were infected with SARS-CoV-2 Omicron variant JN.1, and serially passaged in the presence of either BA.1-D3 (monomer) or BA.1-C2 (trimer). After eight (for BA.1-D3) or nine passages (for BA.1-C2), neutralisation of the passaged viruses was assessed by BA.1-D3 and BA.1-C2 respectively in a micro-neutralisation assay. The results showed that BA.1-D3 had selected for viruses that were no longer neutralised by this nanobody, whereas for viruses passaged in the presence of BA.1-C2, viruses remained susceptible to the nanobody, although with an approximately seven-fold higher NT50 (0.07 nM vs 0.4 8nM) (Fig. 4b, c). In the cross-over experiment BA.1-C2 showed no loss of neutralization potency against the BA.1-D3 escape mutant and vice versa (Fig. 4d, e). This suggested that any sequence changes in the escape variants were independent.

**Figure 4.**
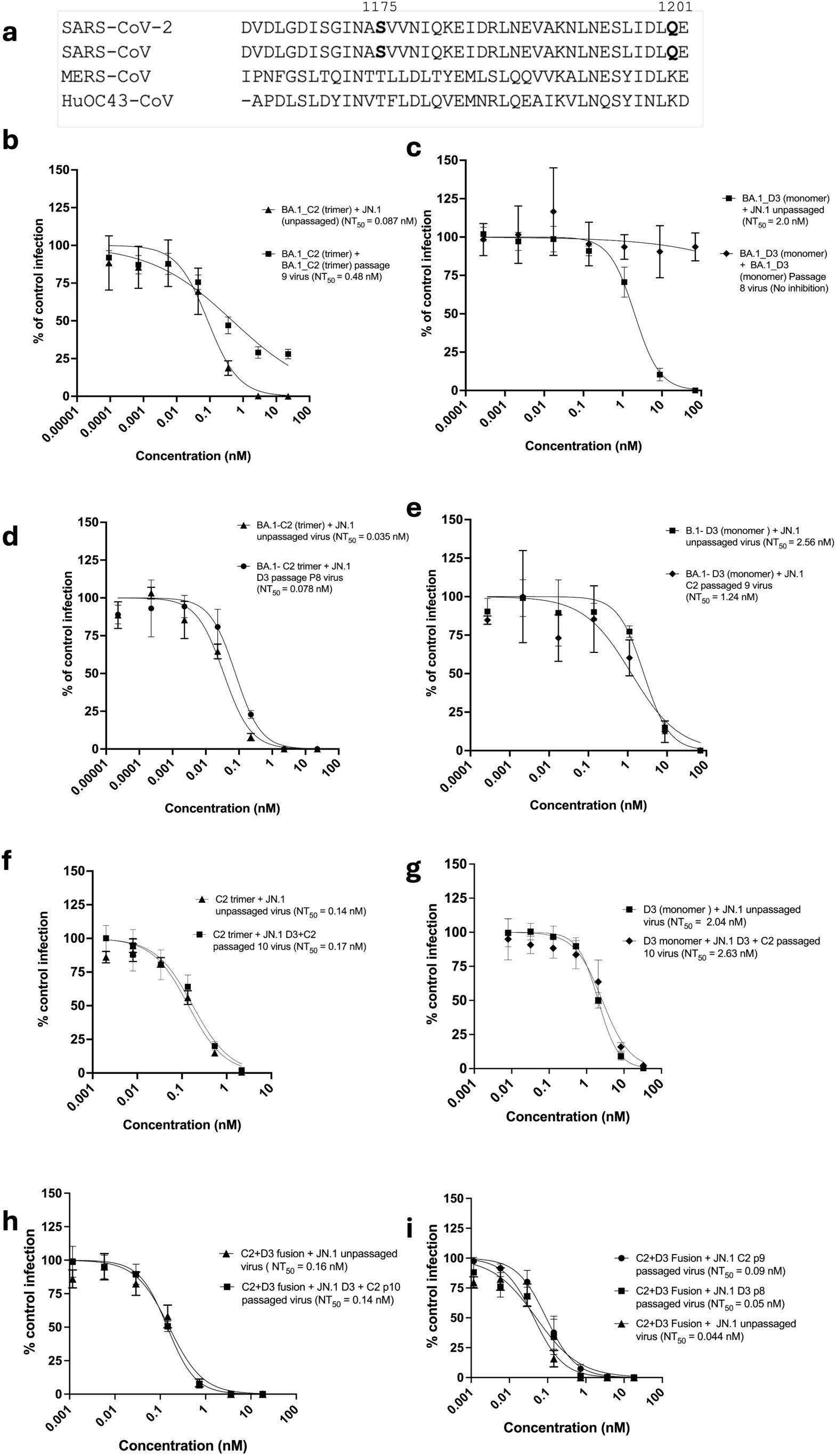
SARS-CoV-2 Omicron JN.1 viral escape. **a** Multiple sequence alignment of the HR2 regions of the spike proteins of HuCoV-OC43, MERS-CoV and SARS-CoV-2 and SARS-CoV (Uniprot ids P36334, R9UQ53, PODCT2, P59594) with the SARS-CoV-2 escape mutants labelled. Neutralisation curves of SARS-CoV-2 Omicron JN.1 by either **b** BA.1-C2 trimer or **c** BA.1-D3 monomer before and after nine passages of the virus in the presence of the corresponding nanobody and **d** by BA.1-C2 before or after eight passages with BA.1-D3, **e** by BA.1-D3 before or after nine passages with BA.1-C2. Neutralization curves of Omicron JN.1 virus before and after ten passages with the BA.1-C2+D3 by **f** BA.1-C2, **g** BA.1-D3 and **h** BA.1-C2+D3 dimer and **i** neutralisation of SARS-CoV-2 by BA.1-C2+D3 before and after either nine passages of the virus in the presence of BA.1-C2 (Q1210K mutant) or eight passages with BA.1-D3 (S1175P mutant). Data are shown as mean (n =2) +/- 95% Cl.

Sequencing of the spike proteins of the escape mutants showed that one mutation (S1175P) in the S2 region of the spike protein conferred resistance to neutralisation by BA.1-D3, and one residue change led to a reduced activity of BA.1-C2 (Q1201K). Both amino acid changes are in the membrane proximal second heptad repeat (HR2) of the spike protein. This sequence is highly conserved between all known variants of SARS-CoV-2 and is identical in SARS-CoV but not MERS-CoV or HuOC43-CoV (Fig. 4a) which accounts for the lack of binding to the spike proteins of these viruses (Fig. 1c). Most interestingly, from alignment of the 1425 full length spike protein sequences with greater than or equal to 50 % sequence identify to SARS-CoV-2 that are returned from a Blast search of Genbank, 94 % have identical HR2 sequences. HR2 forms a triple helix at the base of the spike trimer, and the mutated residues are located at opposite ends of the helices, confirming the experimental result that the epitopes for the BA.1-C2 and D3 nanobodies do not overlap. The D3 escape mutation (S1175P) would prevent N-glycosylation at N1173 by disrupting the NXS/T acceptor sequence [28].

Systematic mutagenesis of N-glycosylation sites in S2 indicates that unlike at other positions, glycosylation at N1173 is not important for spike protein expression or cell-binding [29].

Given that the epitopes of BA.1-C2 and D3 are distinct a second escape mutant study was carried out in which the SARS-CoV-2 Omicron JN.1 virus was passaged in the presence of an equimolar mixture of the two nanobodies as described above. After ten passages, the susceptibility of the virus to neutralization by the either BA.1-C2, BA.1-D3 was tested in a microneutralization assay. The results showed no reduction in the potency of the nanobodies against the virus passaged with an equimolar mixture of the two nanobodies compared to the un-passaged virus (Figure 4 f, g).

A heterodimer was constructed by genetically joining BA.1-C2 and BA.1-D3 with a six residue GlySer linker and the expressed protein assayed for antigen-binding and virus neutralisation activity. The affinity of the BA.1-C2:D3 dimer was similar to the BA.1-D3 monomer showing that the binding is determined by the higher affinity nanobody in the dimer (Supplementary Fig. 1). The neutralisation potency of the dimer was also the same as the BA.1-C2 monomer and homotrimer consistent with the binding affinity (Fig. 4h). The BA.1-C2:D3 dimer was then tested against the escape mutant viruses selected by serial passaging with either nanobody BA.1-C2(Q1201K) or BA.1-D3 (S1175P). The results showed that infection by both mutant viruses was blocked as effectively as the wild-type virus (Fig. 4i). Collectively, the virus escape studies have localised the binding region of the two neutralising nanobodies to the HR2 of the spike protein and shown that the nanobodies combined, either as a mixture or as single heterodimer eliminated viral escape.

### Modelling of BA.1-C2 and BA.1-D3 nanobody-HR2 complexes using Alphafold3

De Cae et al [18] have also localised a neutralising nanobody to the HR2 region of SARS-CoV-2 with Q1201 identified as a key contact residue with the nanobody (R3DC23) indicating an overlap between the epitopes of R3DC23 and BA.1-C2. Structural analysis of the HR2-R3DC23 complex reported by de Cae et al. [18] showed that the epitope of this nanobody involved residues from adjacent helices. To test whether the epitopes of BA.1-C2 and BA.1-D3 overlapped with R3DC23 a competition BLI experiment was carried out with the stabilised BA.1 S2 trimer as the ligand. The results showed that R3DC23 competes with BA.1-C2 but not BA.1-D3 for binding to the S2 trimer and hence the epitopes of R3DC23 and BA.1-C2 must overlap (Supplementary Fig. 2).

The HR2 region is not observed in any of the published cryo-EM structures of isolated spike trimers, even though the sequence is part of the proteins analysed. This presumably reflects the conformational flexibility of the linker between the two heptad repeats and explains why our attempts to determine the structure of BA.1-C2 in complex with a BA.1 spike trimer by cryo-EM were unsuccessful. Therefore, we used Alphafold3 [30] to generate a model of BA.1-C2 in complex with the HR2 which predicted that three nanobodies bound to an HR2 trimer and interacted with residues in adjacent helices (Fig. 5a). AlphaFold produced a model with pTM and IpTM scores of 0.75 and 0.69 respectively. Barring 8 residues at the N - termini, and 6 residues of the C - termini of each HR2 helix, mean pLDDT scores for all residues were >70 (Supplementary Fig. 3). At the interface, per-atom pLDDT scores ranged 57.14 [C2/Arg30/NH2] to 95.08 [C2/Arg99/N], with a mean of 82.34. Nanobody BA.1-C2 was predicted to make an extensive hydrogen bonding network involving the escape-critical Q1201, and side chains of both R101 and N102 of the BA.1-C2 CDR3, as well as N1194, I1198 and E1202 of the same HR2 helix (Chain A) that presents Q1201 to BA.1-C2 (Fig. 5b). The nanobody was further predicted to engage with the adjacent HR2 helix (chain B), via residues of all CDRs (Fig. 5c). Of CDR1, R30 makes a salt bridge with D1199, and R56 of CDR2 makes a salt bridge with E1202. R99 of the CDR3 is predicted to interact with E1188 and N1192, and finally the BA.1-C2 CDR3 backbone provides several additional hydrogen bonds (the amides of I105/N and V104/N bind to the carboxamide of N1192, and the carboxyl of N102 hydrogen bonds to the hydroxyl of S1196).Overall BA.1-C2 is predicted to bind in a very similar location to R3DC23 in the complex with HR2 determined by X-ray crystallography by De Cae et al. [18] consistent with the BLI competition assay results (Supplementary Fig. 2)

**Figure 5.**
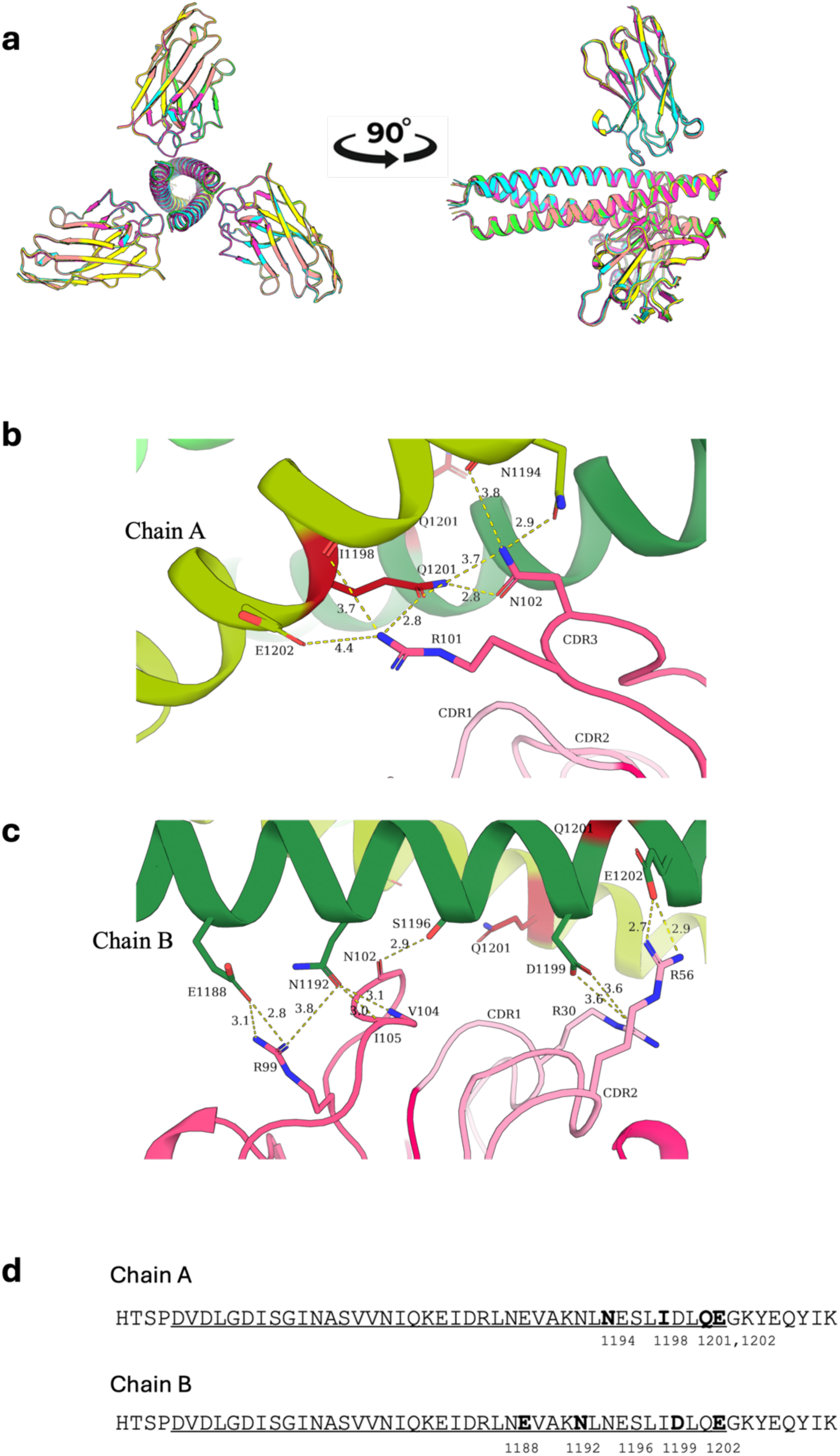
Modelling BA.1-C2: HR2 complex using Alphafold3. **a** Superimposition of top five scoring Alphafold 3 models of the BA.1-C2:HR2 complex. Details of the interface between BA.1-C2 and the helices of **b** Chain A (light green) and **c** Chain B (dark green) hydrogen bonds and salt bridges shown in dotted yellow lines with atomic distances in angstroms (Å) and the CDR1 and 2 loops of BA.1-C2 shown in light pink and CDR3 in dark pink. Figures were generated using The PyMOL Molecular Graphics System, Version 2.5.5 Schrödinger, LLC.

Attempts to model the BA.1-D3: HR2 complex were not successful, generating structures with low confidence scores that did not incorporate the interaction with the S2 escape mutant, S1175P. The models also placed the nanobody in the same location as BA.1-C2 which is not consistent with the results of epitope binning that showed BA.1-D3 does not compete with BA.1-C2 binding to S2.

### BA.1-C2 inhibits cell fusion

Given the location of the binding site of BA.1-C2, a cell-based assay was used to test whether the nanobody inhibited spike-dependent cell fusion. HEK-293T cells were transfected with a vector encoding full length JN.1.11 spike protein and localization of the protein at the cell surface confirmed by immunofluorescence using Alexa Fluro 568-labelled BA.1-C2 or MERS-C9 nanobody (Supplementary Fig.4). Cells expressing the spike protein were incubated with either un-labelled BA.1-C2 or MERS-C9 and added to Calu3 cells stably expressing ACE-2-eGFP [26]. The cells were imaged and the number and size of syncitia quantified. The results showed that BA.1-C2 dose-dependently inhibited cell fusion whereas the negative control MERS-C9 had no effect on syncytial formation (Fig. 6). We conclude that binding of BA.1-C2 to the HR2 of the spike protein blocks cell infection by interfering with fusion events presumably by preventing exposure of the fusion peptide through interfering with the rearrangement of the helices that form the Heptad repeats.

**Figure 6.**
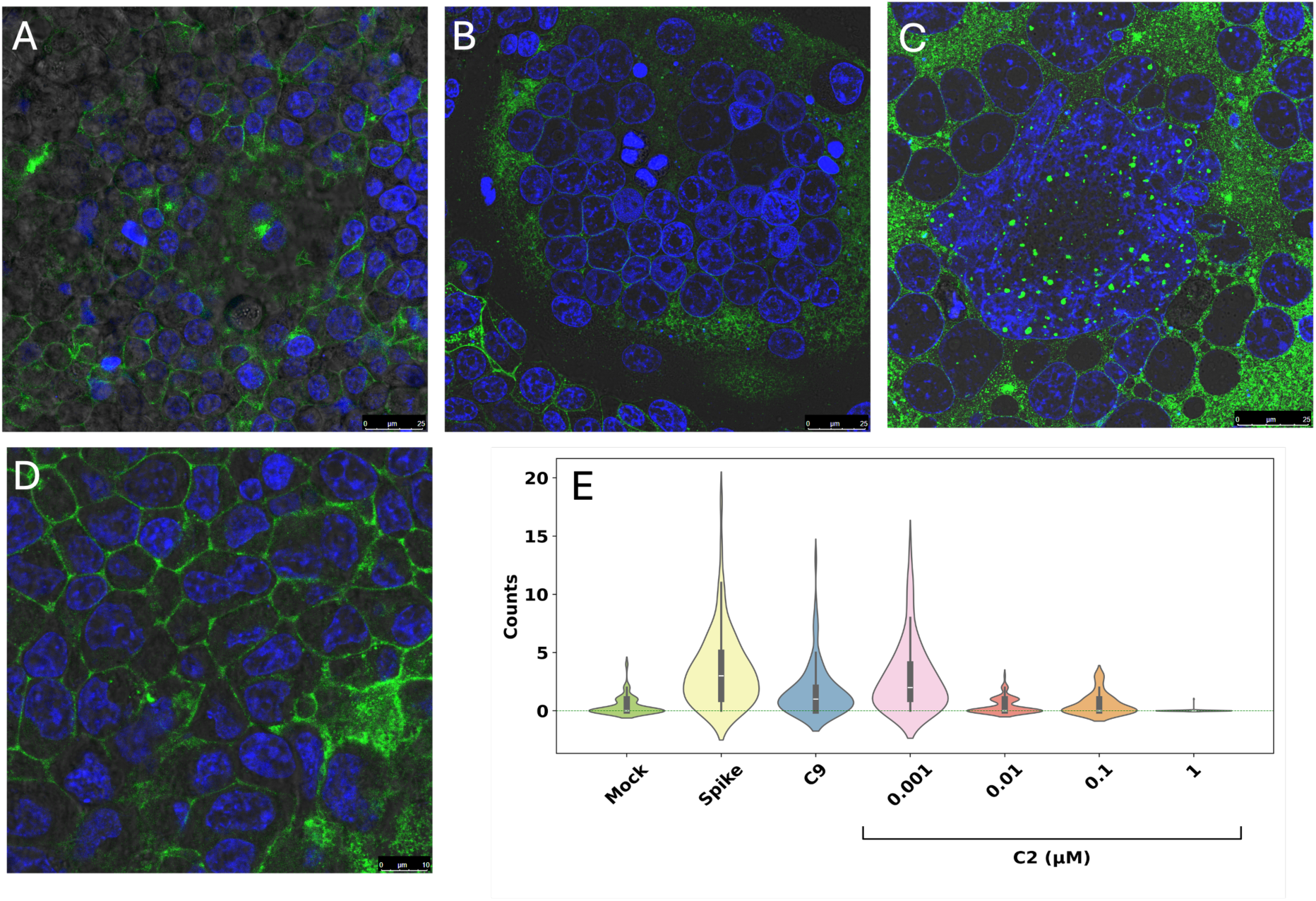
Fusion assay. HEK293T cells expressing JN 1.11.1 spike were incubated with nanobodies for 15 minutes and co-cultured with Calu-3 cells stably expressing ACE2-eGFP to assess syncytia formation. Nuclei were stained with Hoescht 33342 (blue) and ACE2-eGFP shown in green. **a** Non-transfected HEK293T co-cultured with Calu3-ACE2-eGFP cells, **b** HEK293T-JN 1.11.1 spike with Calu3-ACE2-eGFP cells without nanobody treatment, **c** with nanobody MERS-C9, **d** with nanobody BA.1-C2, **e** Violin plots of the number and size of syncytia formed following nanobody treatments: 1µM for MERs-C9, and 1, 0.1, 0.01, 0.001µM for BA.1-C2.

### BA.1-C2 timer provides preventive protection against SARS-CoV-2 Omicron infection of Syrian Golden hamsters

To test whether the neutralization activity observed *in vitro* from targeting the S2 region and to compare with previous results for anti-S1 nanobodies, the efficacy of the BA.1-C2 trimer was evaluated in the hamster model of COVID-19. The study design is shown in Figure 7a, in which BA.1-C2 trimers were administered either 2 h or 24 h before, or 24 h after challenge with SARS-CoV-2 Omicron BA.5. Omicron variants show reduced pathogenicity in the Syrian hamster model [31] compared to earlier strains, reflected by a more modest weight loss in infected animals of 5% *vs* 15-20% observed in a previous study [32] (Fig. 7b). Nonetheless, either prophylactic or therapeutic treatment with the nanobody trimers (2 mg/kg) delivered either intranasally (IN) or into the peritoneum (IP) prevented or reduced weight loss 7 days following challenge with SARS-CoV-2 Omicron BA.5. The most effective treatment regime was intranasal administration 2 h before challenge. Viral RNA load was assessed in throat swabs, nasal and lung tissues by qRT-PCR and whilst no statistically significant differences were observed between throat swabs between treatment and control groups at day 5, by day 7 viral loads in the nasal and lung tissue from all groups was significantly reduced compared to the untreated control. (Fig. 7c and Supplementary Fig. 5) In all treated groups, the extent of parenchymal consolidation was reduced as quantified by automated morphometric analysis which resulted in a statistically significantly larger area of ventilated lung parenchyma compared to the PBS control (Fig. 7d). The results of histology examination of the lungs of control and treated animals are shown in Supplementary Fig. 6.

**Figure 7.**
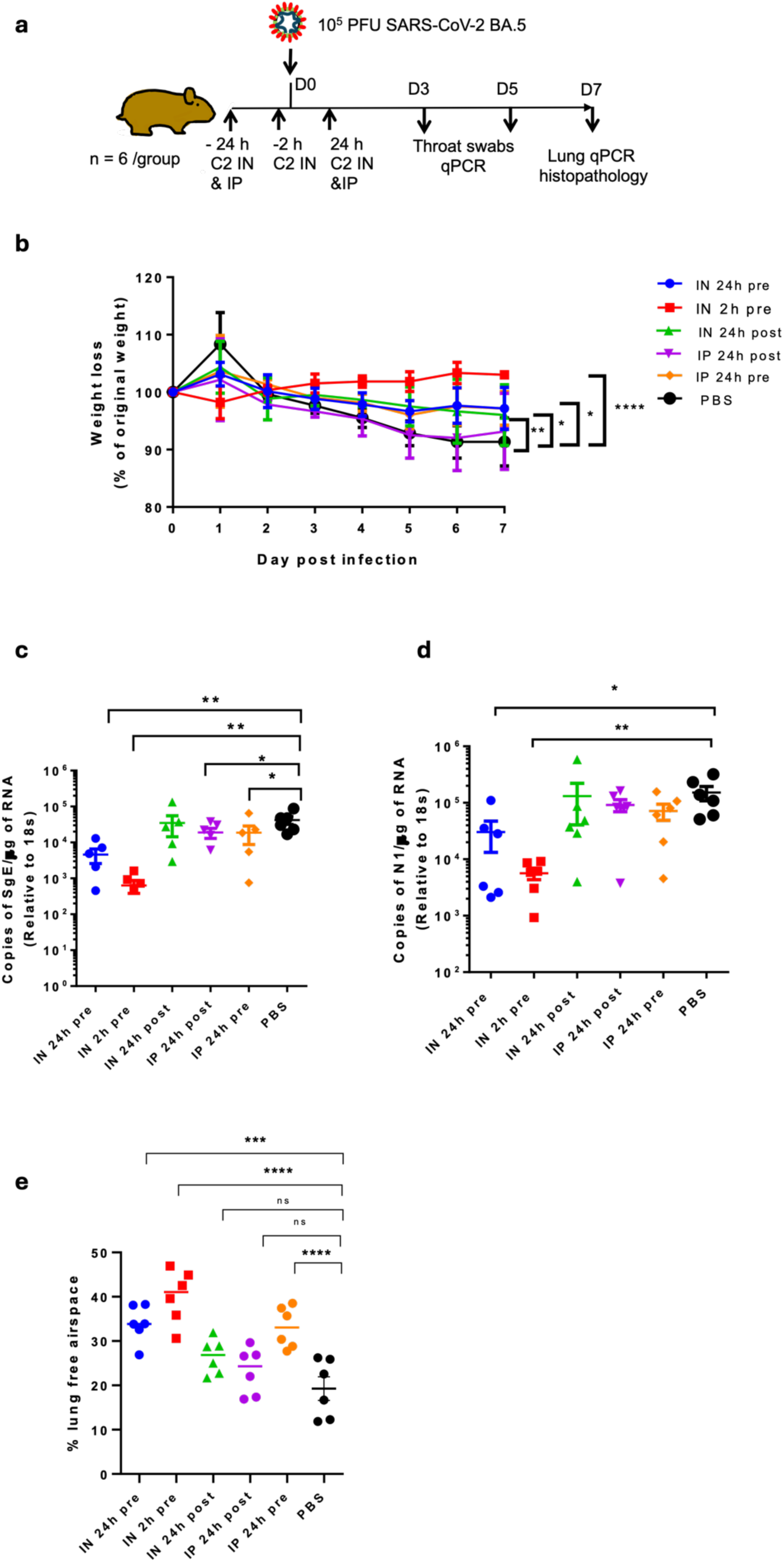
Prophylactic and therapeutic efficacy of BA.1-C2 nanobody trimer in the Syrian hamster model of COVID-19. Golden Syrian hamsters (n = 6 per group) were infected intranasally with SARS-CoV-2 strain Omicron BA.5 (10^5^ pfu in 100 µl PBS). Individual cohorts were treated either 2 h pre-infection, 24 h pre-infection or 24 h post-infection (hpi) with 2 mg/kg 100μl of BA.1-C2 either intranasally (IN) or interperitoneally (IP) as indicated or with PBS alone. **b** Animals were monitored for weight loss at indicated time-points. Data represent the mean value ± SEM. **c** RNA extracted from lung tissue was analysed for SARS-CoV-2 viral load using qRT-PCR for the N gene levels by qRT-PCR and sub-genomic RNA. Assays were normalised relative to levels of 18S RNA. Data for individual animals are shown with the median value represented by a horizontal line. Pairwise comparisons were made between groups using a Mann-Whitney U test. ** represents p < 0.01 and * p < 0.1. **d** Morphometric analysis. HE-stained sections were scanned and analysed using the software programme Visiopharm to quantify the area of non-aerated parenchyma and aerated parenchyma in relation to the total area. Results are expressed as the mean free airspace in lung sections. Pairwise comparisons were made between groups using a Mann-Whitney U test. *Represents p < 0.05; ** represents p < 0.01.

## Discussion

The S2 region of SARS-CoV-2 and related beta-coronaviruses comprises several segments, including the fusion peptide, Heptad repeats 1 (HR1) and 2 (HR2) that play key structural and functional roles in virus entry. By targeting the S2 region of the spike proteins of pathogenic SARS-CoV-2 we have identified nanobodies that selectively neutralize the virus *in vitro*.

Further, one of the nanobodies to the S2 of SARS-CoV-2 (BA.1-C2) produced as a homotrimer was shown to reduce lung viral load and associated weight loss in the Syrian golden hamster model of COVID-19. The results are comparable to our previous study in which animals challenged with the same SARS-CoV-2 variant (Omicron BA.5) were treated with nanobodies targeting the S1 RBD [26]. Consistent with the results of De Cae et al [18] we have shown the therapeutic potential of targeting the HR2 segment of the S2 region of the spike protein with a single domain antibody.

The sub-nanomolar neutralization potencies of the S2 binding nanobodies reported are typically greater than conventional antibodies that have been mapped to S2 region (NT50 between 10-50 nM) [5, 7, 14] pointing to recognition of different epitopes or binding modes by nanobodies compared to antibodies. The binding sites of the nanobodies to SARS-CoV-2, BA.1-C2 and BA.1-D3 were localised to the membrane proximal region Heptad repeat 2 (HR2),which is also the binding site of a neutralizing nanobody previously identified [18].

The HR2 sequences of SARS-CoV-2 and SARS-CoV are identical and earlier work had shown that rabbit polyclonal antibodies raised to linear HR2 sequences have virus neutralization activity [33, 34]. The HR2 sequence is not typically a binding site of patient derived monoclonal antibodies to the S2 region spike protein of SARS-CoV-2 with the fusion peptide (residues 809-834) and stem-helix (residues 1140-1168) that only includes the six N-terminal residues of HR2, presenting the immunodominant neutralizing epitopes [4, 5, 9, 17, 35]. However, more recently, a human monoclonal antibody that binds to the HR2 of SARS-CoV-2 has been characterized [36]. The neutralizing nanobodies reported here and elsewhere [18] suggests that the HR2 sequence could be a promising candidate for vaccine development. The interaction between HR1 and HR2 plays a key role in forming the six-helical bundle fusion core that mediates viral fusion and involves the packing of the HR2 segment into grooves formed by an extended HR1 coiled-coil triggered by furin cleavage between S1 and S2 [37]. SARS-CoV-2 HR2 peptides with [38, 39] or without additional HR1 sequences [40–43] have been shown to inhibit virus fusion and the mechanism of action of these inhibitors is presumed to be preventing the formation of the fusion core by disrupting the interaction between HR1-HR2. It seems likely that the nanobodies which bind to HR2 work in a similar way. The mutant escape study reported here and by de Cae et al [18] showed the potential for specific residue changes in the HR2 to abrogate or reduce nanobody binding, notably, S1175, Q1201 (this study) and N1192,L1197, L1200 and also Q1201 [18]. Profiling of SARS-CoV-2 escape mutants for Spike-binding antibodies using phage-deep mutational scanning revealed that for the convalescent patient sera that were screened, sensitivity to mutations at different sites in linker/HR2 region was highly variable [35]. We showed that combining nanobodies BA.1-C2 and BA.1-D3 that bind to different epitopes in the HR2 sequence, mitigated the effects of selecting for escape mutations. Therefore, the heterodimer has potential as a prophylactic treatment administered intranasally, that could provide protection for immunocompromised individuals during periods of high disease burden in the community. Given the exceptional level of sequence conservation amongst bat Sarbecoviruses in the HR2 region recognized by these nanobodies, it seems possible that the nanobodies could also be protective against future emergent pathogenic viruses.

## Methods

### Immunization and construction of VHH library

Vectors for expressing SARS-CoV-2 (Omicron BA.1), HuCoV-OC43 and MERS-CoV trimeric spike proteins, containing twin proline substitutions and mutated furin cleavage sites, were generously provided by Piyada Supasa and Gavin Screaton (Nuffield Department of Medicine, University of Oxford, Oxford, UK). Trimeric spike proteins were produced [14], and antibodies were raised in a llama as previously described [1]. Briefly, each spike protein (200 μg) was mixed with the adjuvant Gerbu LQ#3000 for each of the three intramuscular immunizations of the same llama on days 0, 28 and 56. Blood (150 ml) was collected on day 66. Immunizations and handling of the llama were performed under the authority of the project license PA1FB163A. VHHs were amplified by two rounds of PCR from cDNA prepared from peripheral blood monocytes and cloned into the SfiI sites of the phagemid vector pADL-23c (Antibody Design Laboratories, San Diego, CA, USA). Electro-competent *E. coli* TG1 cells (Agilent Technologies LDA UK) were transformed with the recombinant pADL-23c vectors, and the resulting TG1 library stock was infected with M13K07 helper phage to obtain a library of VHH-presenting phages.

### Isolation of VHHs and construction of Fc fusions, VHHs dimers and trimers

Phage displaying VHHs specific for the S2 regions were enriched by two rounds of panning on 50 nM and 5 nM of biotinylated S2 regions, through capturing with Dynabeads M-280 (Thermo Fisher Scientific). After the second round of panning, 93 individual phagemid clones were picked, VHH-displaying phage were recovered by infection with M13K07 helper phage and tested for S2 binding by ELISA with biotin-tagged RBDs immobilized on neutravidin-coated plates. Positive phage binders were sequenced and grouped according to CDR3 sequence identity using the IMGT/V-QUEST server [46]. Heterodimeric and homotrimeric VHH constructs were generated by total gene synthesis (ITD technology) with [GlySer]_3_ linkers connecting the VHHs. The gene products were inserted into the pOPINTTGneo vector by Infusion® cloning. Vectors for expressing VHHs as C-terminal human IgG1 Fc fusions were constructed by PCR amplification of the VHHs and cloning into pOPINTTG-3C-IgG1Fc. Both pOPINTTGneo and pOPINTTG-3C-IgG1Fc contain a mu-phosphatase leader sequence and C-terminal His6 tag [47].

### Protein production

VHH plasmids were transformed into the WK6 *E. coli* strain and protein expression was induced by 1 mM IPTG during overnight growth at 28°C. Periplasmic extracts were prepared by osmotic shock and VHH proteins were purified by immobilised metal affinity chromatography (IMAC) using an automated protocol implemented on an ÄKTXpress followed by gel filtration using a Hiload 16/60 Superdex 75 or a Superdex 75 10/300 GL column, using PBS pH 7.4 buffer. Trimeric versions of the S2 domains of Omicron BA.1 (residues 681 to 1200) MERS-CoV (residues 749 to 1291) with a C-terminal Avitag™ were constructed by PCR amplification using the spike trimers as templates. All amplified S2 DNAs were inserted into the vector, pOPINTTGneo using Infusion® cloning [44].The sequences of the expressed spike and S2 proteins are given in the Supplementary information Table 2. The nanobody dimer, trimers, Fc fusions and spike proteins were produced by transient expression in Expi293F cells and purified by a combination of IMAC and gel filtration in PBS pH 7.4 buffer. For animal studies, the final purified product was passed through two Proteus NoEndo clean-up columns (Generon, Slough, UK) to reduce endotoxin to <0.1 EU ml−1. Endotoxin levels were quantified using the Pierce LAL Chromogenic Endotoxin Quantitation Kit (Thermofisher Scientific).

### ELISA

96-well ELISA plates were coated overnight with 1 µg/well neutravidin in PBS and then washed x 5 with PBS containing Tween 20 (0.05 % v/v) (PBST). Biotinylated S2 trimers (50 nM) were added to each well (100 µl) and allowed to bind for 1 h. at RT on a vibrating shaking platform. Plates were blocked for 1h at RT by adding 2 % milk powder in PBST washed x 5 with PBST, and then tenfold serial dilutions of VHH-Fcs (100 to 0.0001 ug/well) added to the plate. Following further incubation for 1 h. at RT with agitation, plates were washed and bound VHH-Fcs detected by the addition of an anti-human IgGFc-HRP conjugate using ABTS substrate (mix solution A: solution B in a 1:1 ratio). Colour development was measured by absorbance at 405 nm. For epitope mapping, binding of VHH-Fcs was assayed in the presence of Fab B6 (10 µg/ml). Binding of the Fab fragment of B6 was detected using an anti-mouse light chain-HRP conjugate.

### Biolayer interferometry

Biolayer interferometry was used to measure the binding constants of the nanobodies to SARS-CoV-2 spike and S2, and for epitope binning [26]. All assays were performed using a Sartorius Octet R8 system and designed using Octet BLI Discovery 12.2.2.20 software for normalization of the association and dissociation steps and Savitzky-Golay filtering. Curve fitting was applied using a global fit method and the association and dissociation rates calculated using a best fit method. All graphs were plotted using GraphPad Prism.

### Microneutralization assay

SARS-CoV-2 microneutralization assays were carried out as previously described [26]. The neutralization titre (NT_50_) was defined as the titre of either the monomeric, dimeric Fc fusions or trimeric VHHs that reduced the Foci forming unit (FFU) by 50% compared to the control wells.

### Isolation and sequencing of escape mutants

BA.1-C2 (trimer) and BA.1-D3 (monomer) were passaged with SARS-CoV-2 JN.1 at an MOI of 0.005 in Vero E6 TMPRSS2 cell line, initially at the NT_50_ for two passages then at the NT_90_ for passages three to five, followed by doubling the concentration for subsequent passages. Microneutralisation assays were carried out as previously described in between passaging to determine potential reductions in neutralisation. RNA was extracted from supernatant of passage 8 for BA.1-D3 (monomer) and passage 9 for BA.1-C2 (trimer) using Qiagen QIAamp Viral RNA Mini Kit, according to manufacturer’s procedure. RNA was prepared for Illumina sequencing using NEB NEBNext® Ultra™ II FS DNA Library Prep Kit for Illumina®. Raw reads were processed using Galaxy server version 24.2.4. dev0 by mapping the reference genome (GenBank Accession: PP832909.1) and variants called for mutations and associated frequencies. Mutations were verified using NextClade https://clades.nextstrain.org [45]

### AlphaFold Modelling

The interaction of BA.1-C2 with the SARS-CoV-2 HR2 helical bundle was predicted using AlphaFold 3 [30]. Three copies of the SARS-CoV-2 HR2 amino acid sequence (residues 1163-1209), along with three copies of the complete C2 VHH amino acid sequence (contiguous residues 1-126) were submitted to https://alphafoldserver.com, using a prediction seed of 100. Models were visualised using The PyMOL Molecular Graphics System, Version 2.5.5 Schrödinger, LLC.

### Fusion assay

HEK293T cells were maintained in phenol red free DMEM supplemented with 1% GlutaMAX and 10% FBS at 37°C with 5% CO2. Calu-3 cells stably expressing ACE2-eGFP [26] were grown in Phenol red free DMEM-F12 media (Thermo Fisher Ltd.,) supplemented with 1% GlutaMAX™ (Thermo Fisher Ltd.,) and 10% FBS under the same incubation conditions. For experiments, HEK293T cells were seeded in 24-well plates at the density of 1.5 x10^4^ cells per well. After 16 hours, cells were transiently transfected with plasmid DNA encoding JN1.11.1 spike (Addgene plasmid no: 223272, pTwist-SARS-CoV2 Spike-delta C18) following the manufacturer’s protocol. Briefly, 500 ng of JN.1.11.1 Spike-delta, 1µl of Fugene HD (Promega) and 50 µl of serum free DMEM were mixed, incubated for 5 minutes at room temperature and added dropwise to each well. Nanobodies MERs-C9 and BA.1-C2 were fluorescently labelled with Alexafluor 568 via cysteine-maleimide conjugation as described previously [46]. After 24 hrs post-transfection, HEK293 cells were treated with different nanobodies for 15 minutes (1 µM), washed and incubated with fresh media. Cells were imaged by laser scanning confocal microscope with a 60-x oil-immersion objective. (DMi8, Leica Biosystems Ltd., UK) to confirm nanobody binding to the spike protein.

HEK293T cells expressing JN 1.11.1 spike labelled with nanobodies were co-cultured with Calu-3 cells stably expressing ACE2-eGFP by adding 1 × 10⁴ cells per well in a 24-well plate (Ibidi GmbH) in DMEM supplemented with GlutaMAX™ and 10% FBS. After 36 hours, cells were labelled with Hoechst 33342 (1 µg/mL) for 10 minutes, washed twice with PBS, and fixed with 4 % (v/v) paraformaldehyde for 10 minutes at room temperature. Following fixation, cells were washed twice with PBS and prepared for imaging. All wells were imaged using a laser-scanning confocal microscope with a 20-x objective, and 150 images per well were collected. Syncytia formation was quantified across different nanobodies and concentrations. In addition to the quantitative imaging, selected fields were imaged using a 60x oil immersion objective to obtain high-resolution images for presentation.

### Evaluation of trimer prophylactic & therapeutic efficacy in the Syrian Golden hamster model

Animal work was approved by the local University of Liverpool Animal Welfare and Ethical Review Body and performed under UK Home Office Project License PP4715265. Male Syrian Golden hamsters (8-10 weeks old) were purchased from Janvier Labs (France). Animals were maintained under SPF barrier conditions in individually ventilated cages. For virus infection, an Omicron BA.5 strain (MTA free) of SARS-CoV-2 was used, kindly provided by Prof Wendy Barclay (Imperial college, London), isolated from the UK in June 2022 and sequence verified. Animals were randomly assigned into multiple cohorts of 6 animals. For SARS-CoV-2 infection, hamsters were anaesthetised lightly with isoflurane and inoculated intra-nasally with 100 µl containing 10^5^ PFU SARS-CoV-2 in PBS. Hamsters were treated either via the intranasal or inter-peritoneal routes with 100 µl BA.1-C2 trimers in PBS (2 mg/kg) at either 2h or 24 h before virus challenge or 24 h post-infection. Throat swabs were taken on day 3 and day 5 post-challenge and animals sacrificed on day 7 by an overdose of pentobarbitone. RNA was isolated from oropharyngeal swabs and tissue samples (lung and nasal turbinate tissue). Right upper lobe of lung and nasal tissue samples were homogenized and inactivated in 1 ml of Trizol reagent (Thermofisher) using stainless steel bead and a tissue lyser LT (Qiagen) at 50 hz oscillation for 5 minutes. Oropharyngeal swabs samples were inactivated in Trizol LS at 1:3 (Invitrogen) Tissue homogenate was then centrifuged at 12000xg for 5 min and RNA extraction was performed as per manufacturer’s instructions. Downstream extraction was then performed using the BioSprintTM96 One-For-All vet kit (Indical Bioscience) and Kingfisher Flex platform as per manufacturer’s instructions. Non-tissue samples were inactivated in AVL (Qiagen) and ethanol, with final extraction using the BioSprintTM96 One-For-All vet kit (Indical Bioscience) and Kingfisher Flex platform as per manufacturer’s instructions. Reverse transcription-quantitative polymerase chain reaction (RT-qPCR) was performed using TaqPathTM 1-Step RT-qPCR Master Mix, CG (Applied BiosystemsTM), 2019-nCoV CDC RUO Kit (Integrated DNA Technologies) and Bio-Rad CFX mastero Real-Time PCR System. Sequences of the N1 primers and probe were: 2019-nCoV_N1-forward, 5’ GACCCCAAAATCAGCGAAAT3’;2019-nCoV_N1-reverse,5’TCTGGTTACTGCCAGTTG AATCTG 3’; 2019-nCoV_N1-probe, 5’ FAM-ACCCCGCATTACGTTTGGTGGACC-BHQ1 3’. The cycling conditions were 25°C for 2 min, 50°C for 15 min, 95°C for 2 min, followed by 45 cycles of 95°C for 3 seconds, 55°C for 30 seconds. The quantification standard was in vitro transcribed RNA of the SARS-CoV-2 N ORF (accession number NC_045512.2) with quantification between 10 and 1x106 copies/μl. Positive samples detected below the lower limit of quantification (LLOQ) of 10 copies/μl were assigned the value of 5 copies/μl, undetected samples were assigned the value of 2.3 copies/μl, equivalent to the assays LLOD. For nasal wash and oropharyngeal swab extracted samples this equates to an LLOQ of 1.29 x104 copies/mL and LLOD of 2.96 x103 copies/mL. Samples detected between LLOQ and LLOD were assigned 6.43 x103 copies/mL. For tissue samples this equates to an LLOQ of 1.31x104 copies/g and LLOD of 5.71 x104 copies/g. Samples detected between LLOQ and LLOD were assigned 2.86 x104 copies/g. Subgenomic RT-qPCR was performed on the QuantStudioTM 7 Flex Real-Time PCR System using TaqManTM Fast Virus 1-Step Master Mix (Thermo Fisher Scientific) and oligonucleotides as specified by Wolfel et al67., with forward primer, probe and reverse primer at a final concentration of 250 nM, 125 nM and 500 nM respectively. Sequences of the sgE primers and probe were: 2019-nCoV_sgE-forward,5’CGATCTCTTGTAGATCTGTTCTC3’;2019-nCoV_sgE reverse,5’ATATTGCAGCAGTAC GCACACA 3’; 2019-nCoV_sgE-probe, 5’ FAM-ACACTAGCCATCCTTACTGCGCTTCG-BHQ1 3’. Cycling conditions were 50°C for 10 minutes, 95°C for 2 min, followed by 45 cycles of 95°C for 10 seconds and 60°C for 30 seconds. RT-qPCR amplicons were quantified against an in vitro transcribed RNA standard of the full-length SARS-CoV-2 E ORF (accession number NC_045512.2) preceded by the UTR leader sequence and putative E gene transcription regulatory sequence described by Wolfel et al. in 202049. Positive samples detected below the lower limit of quantification (LLOQ) were assigned the value of 5 copies/μl, whilst undetected samples were assigned the 29 values of ≤0.9 copies/μl, equivalent to the lower limit of detection of the assay (LLOD). For nasal washes and oropharyngeal swabs extracted samples this equated to an LLOQ of 1.29x104 copies/mL and LLOD of 1.16x103 copies/mL. For tissue samples this equates to an LLOQ of 5.71x104 copies/g and LLOD of 5.14x103 copies/g.

Tissues were removed immediately for downstream processing. From all animals the left lung was fixed in 10% buffered formalin for 48 h and then stored in 70% ethanol until further processing. Two longitudinal sections were prepared and routinely paraffin wax embedded. Consecutive sections (3-5 µm) were prepared and stained with hematoxylin eosin (HE) for histological examination or subjected to immunohistological staining. Immunohistology was performed to detect SARS-CoV-2 antigen, using the horseradish peroxidase (HRP) method and the following primary antibody: rabbit anti-SARS-CoV nucleocapsid protein (Rockland, 200-402-A50) as previously described [32]. For morphometric analysis, the HE-stained sections were scanned (NanoZoomer-2.0-HT; Hamamatsu, Hamamatsu City, Japan) and analysed using the software programme Visiopharm (Visiopharm 2020.08.1.8403; Visiopharm, Hoersholm, Denmark) to quantify the area of non-aerated parenchyma and aerated parenchyma in relation to the total area (= area occupied by lung parenchyma on two sections prepared from the left lung lobes) in the sections, as previously described [32].

## Acknowledgements

This work was supported by the Rosalind Franklin Institute, funding delivery partner EPSRC, Wellcome Trust (223733/Z/21/Z), BBSRC (BB/V018523/1) grants. We thank the laboratory staff of the Histology Laboratory, Institute of Veterinary Pathology, Vetsuisse Faculty, University of Zurich, for excellent technical support, and Professor Gary Stephens, Barney Jones, Hong Lin (Reading University) for expertise in llama immunization.

## Author contributions

J.C., I.B., K.C. H.C. L.McC & L.E. identified/characterised the nanobodies and produced proteins for the experiments. J.C. generated and analysed the Alphfold3 models. L.J. and E.P. carried out and analysed the neutralization assays and virus escape studies. P.S., D.M. and A.K. carried out the animal study A. Kipar performed pathological analyses, immunohistology and morphometric analyses. SR and A. Karmakar carried out and analysed the cell fusion assays, R.J.O, M.C. J.P.S., directed the studies. R.J.O., M.C. W.J. and J.P.S planned the project interpreted the data and wrote the manuscript with contributions from all authors.

