## Supplementary information for "Nanobodies against the S2 region of the spike protein potently neutralize SARS-CoV-2 viruses and show resistance to virus escape"

Supplementary Fig. 1. Binding affinity of C2-D3 fusion protein measured by BLI

Supplementary Figure 2. Competition between BA.1-C2, BA.1-D3 and R3DC23 for binding to BA.1S2 trimer measured by BLI

Supplementary Figure 3 AlphaFold pLDDT scores figure

Supplementary Figure 4. HEK293T cells expressing the JN.1.11.1 spike protein on the cell surface

Supplementary Figure. 5 Viral load in nasal swabs at day 5 and nasal tissues at day 7

Supplementary Figure. 6 Lung histology

Supplementary Table 1 VHH DNA sequences (nos. 1-13 from BA.1 S2 panning only; nos. 14-18 from BA.1 S2 and MERS S2 panning and nos. 19-24 from MERS S2 panning only)

Supplementary Table 2. Sequences of beta-coronavirus spike and S2 proteins

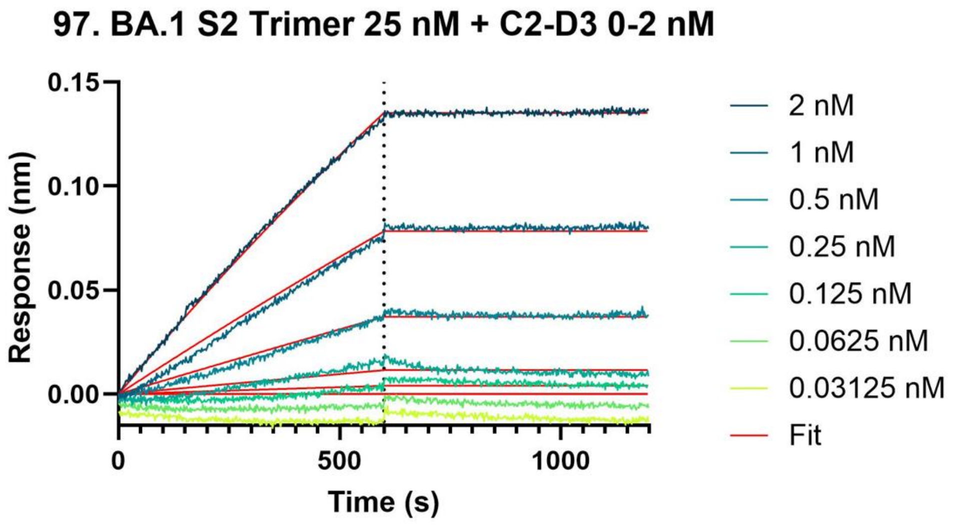

KD: $1.175 \times 10^{-12} \pm 7.070 \times 10^{-13}$ M
$K_a$: $1.192 \times 10^{5} \pm 4.032 \times 10^{3}$ 1/Ms
$K_d$: $1.522 \times 10^{-7} \pm 7.070 \times 10^{-8}$ 1/s

**Fig. 1. Binding affinity of C2-D3 fusion protein measured by BLI**
Biotin-tagged BA.1 S2 trimer (25 nM) was loaded onto streptavidin sensors and then the BA.1- C2-D3 added (0 - 2 nM)

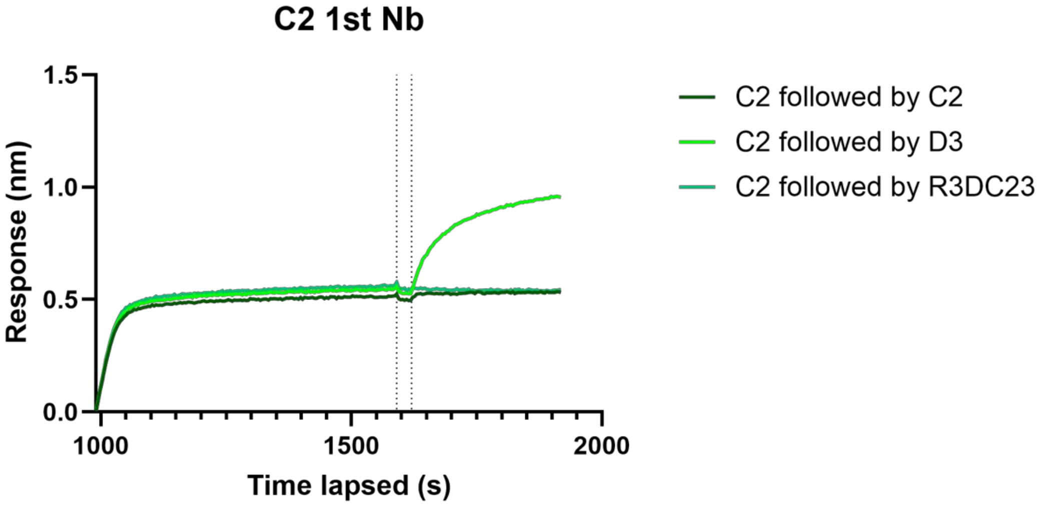

**Fig. 2 Competition between BA.1-C2, BA.1-D3 and R3DC23 for binding to BA.1S2 trimer measured by BLI**
Biotin-tagged BA.1 S2 trimer (200nM)was loaded onto streptavidin sensors and then the BA.1-C2 added (50 nM) followed by either BA.1-C2 (50 nM), BA.1-D3 (50nM) or R3DC23 (100 nM)

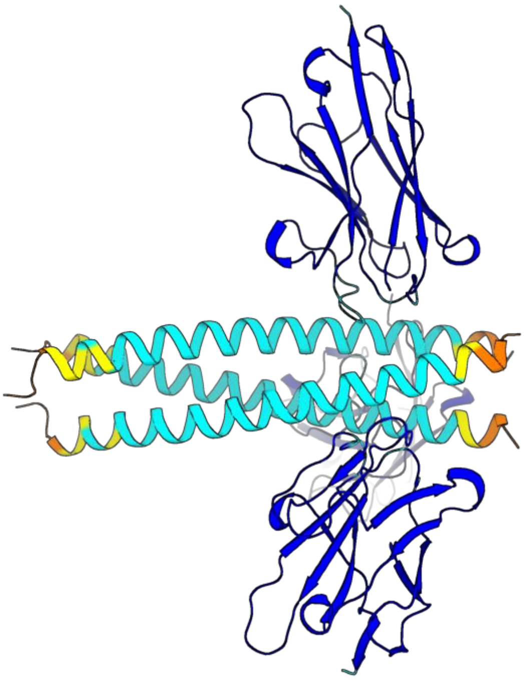

**Fig. 3 AlphaFold pLDDT scores figure**
Ribbon notation for rank 0 HR2-C2 model from AlphaFold Predictions, coloured following standard AlphaFold pLDDT scale: >90; blue, >70; light blue, >50; yellow, <50; orange.

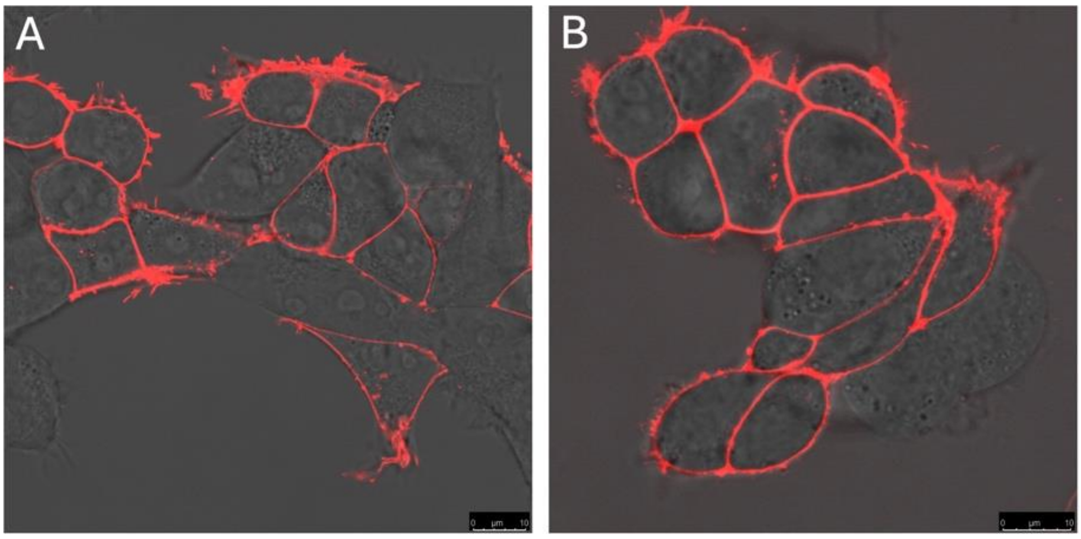

**Fig. 4 HEK293T cells expressing the JN.1.11.1 spike protein on the cell surface**
Nanobodies (A) BA.1- C2, and (B) MERS-C9 were conjugated to Alexa Fluor 568 and incubated with the cells at a final concentration of 1 µM in fresh culture medium for 15 minutes under standard cell culture conditions. Following incubation, the nanobody- containing media were replaced

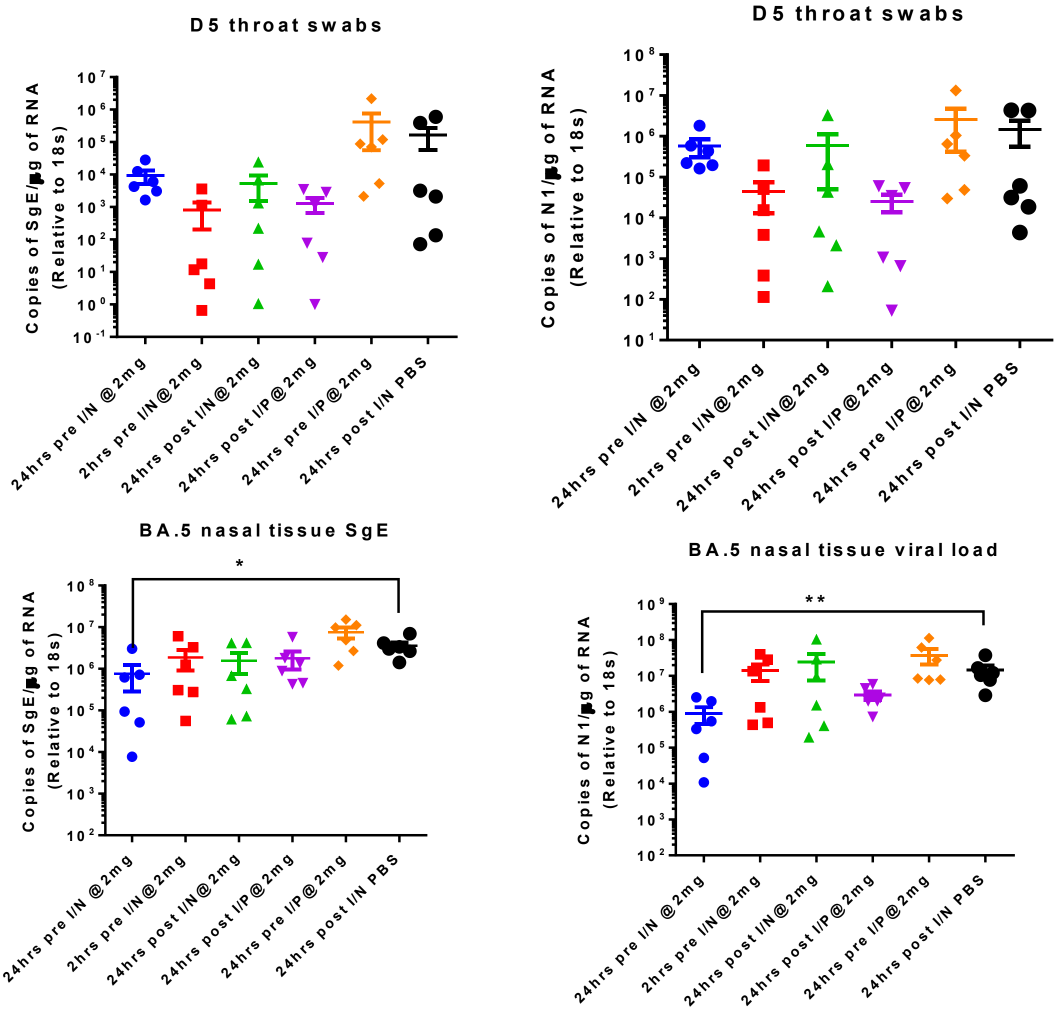

**Fig. 5 Viral load in nasal swabs at day 5 and nasal tissues at day 7**

RNA was analysed for SARS-CoV-2 viral load using qRT-PCR for the N gene and sub-genomic RNA levels by qRT-PCR.Assays were normalised relative to levels of 18S RNA. Data for individual animals are shown with the median value represented by a horizontal line. Pairwise comparisons were made between groups using a Mann-Whitney U test. ** represents $p < 0.01$ and * $p < 0.1$.

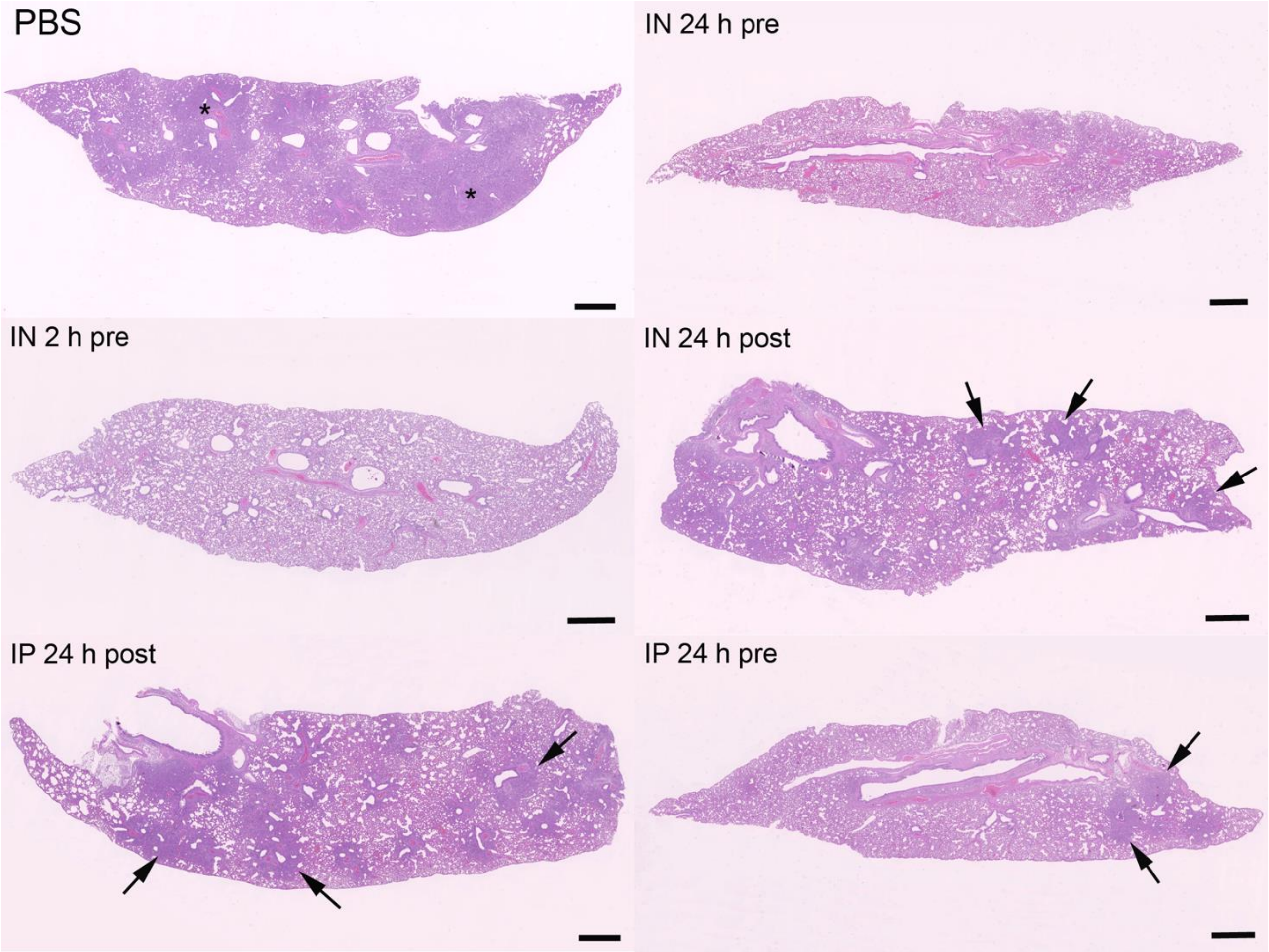

**Fig. 6 Lung histology**

Extent of inflammatory processes in the lungs of hamsters challenged intranasally with SARS-CoV-2 Omicron BA.5 at $10^5$ PFU that had received C2 trimer intranasally (IN) or intraperitoneally (IP) at 24 h or 2 h pre or post infection, or PBS, and were examined 7 days post infection. HE stain, Bars = 1 mm. In control animals (**PBS**), large areas of the lung parenchyma are consolidated (asterisks), with abundant activated and hyperplastic type II cells and multiple areas of type II pneumocyte/bronchial epithelial cell hyperplasia, accompanied by infiltrating macrophages, lymphocytes and neutrophils as well as desquamed alveolar macrophages/type II pneumocytes. Animals that had received C2 trimer intranasally at 24 h and at 2 h prior to infection (**IN 24 h pre, IN 2 h pre**), the lungs do not exhibit any histological changes. When C2 trimer had been applied 24 h post infection, intranasally or intraperitoneally (**IN 24 h post, IP 24 h post**), the lungs exhibit random small consolidated areas (arrows) of a similar composition as control animals (PBS). With intranasal application of C2 trimer at 24 h prior to infection (**IP 24 h pre**), a few small consolidated areas (arrows) of a similar composition as control animals (PBS) are observed.

**Supplementary Table 1** VHH DNA sequences (nos. 1-13 from BA.1 S2 panning only; nos. 14-18 from BA.1 S2 and MERS S2 panning and nos. 19-24 from MERS S2 panning only)

| Number | VHH id | DNA sequence |
|---|---|---|
| 1 | BA.1_D12 | CAGGTGCAGCTGCAGGAGTCTGGGGGAGGATTGGTGCAGCCTGGGGACTCTCTGACACTCTCCTGTGCAGCCTCTGGGCGCGGCTTCGATGCCTACGGCATGATGTGGTTCCGCCAGGCTCCAGGGAAGGAACGTGAATTTGTAACAGGAATTAACTGGGGTGGTACTACATACTATGCGGACTCCGCGAAGGGCCGATTCACCATCTCCAGAACCAGTGACAAGACCACGGTATATCTACAAATGACCAGCCTGAAACCTGAGGACTCGGCCATTTATTACTGTGCAGCAGCTCCCACCAGCACAGTGGTGACTAGACCCTCTCGTCCGAAGTACTGGGGCCAGGGGACCCAGGTCACCGTCTCCTCG |
| 2 | BA.1_A7 | CAGGTGCAGCTGCAGGAGTCTGGGGGAGGATCGGTGCAGGCTGGGGGCTCTCTGAGACTCTCCTGTGTAGCCTCCGGACGCACCGTCAGTCTTTTTGACATGGGCTGGTTCCGCCAGGCTCCAGGGAAGGAGCGTGAATTTGTAGCGCGTATTACGTTGAAGGAAGGTAAACTAAACTATGCAGACTCCGTGAAGGGCCGATTCACCATCTCCAGAGGCAACCCCGAGAACACGGTGTATCTGCAGATGGATAGTCTGAAACCGGAGGACACGGCCGTTTATTACTGTGCAGCAGACCAAACGGTAGTACGTATGACTGGGACCGGGATCGACTACTGGGGCCAGGGGACCCAGGTCACCGTCTCCTCA |
| 3 | BA.1_D2 | CAGGTGCAGCTGCAGGAGTCTGGGGGTGGCGTGGTGCAGGCCGGGGGCTCTCTGAGACTCTCCTGTGCAGCCTCTGGACGCGCCTTCAGCGTCACTACTGTGGCCTGGTTCCGCCAGTCTCCAGGGAAGGAGCGTGAGTACATAGCACGCTCCGACGCCAGAGGCGGTAAATATTATACAGACTCCGTGAAGGGCCGATTCACCATCTCCGACGAACGTGACAAGATGACAGTATATCTACAAATGAACGACCTCGAAACTAGCGACACGGCCGTTTATTATTGTGCAGCCGGCCCGTTTGGAGTCAGCGCTAGAGAAGATGACTATGCTTACTGGGGGCAGGGGACCCAGGTCACCGTCTCCTCA |
| 4 | BA.1 _A5 | CAGCTGCAGGAGTCTGGGGGAGGATTGGTGCAGGCTGGGGGCTCTCTGAGACTCTCCTGTGCCACCTCTGGACGCATTCTCAGTAACTATGTGATGGGCTGGTTCCGCCAGGCTCCAGGGAAGGAGCGTGCGTTACTTGGAGCTATTACCTGGAGCGCAGGTAGAACAGCCTATGCGGAGTCCGTGAAGGGCCGATTCACCATCTCCAGAGACATCGCCGAGAACGCGGTGTATCTGCAAATGAACAGCCTGAAACCTGAGGACACGGCCGTTTATTACTGTGCGGCACGAATTGTATCCGCATCTGAATATACCTACTGGGGCCAGGGGACCCAGGTCACCGTCTCCTCA |
| 5 | BA.1_B10 | CAGGTGCAGCTGCAGGAGTCTGGGGGAGGATTGGTGCAGGCTGGAGGCTCTCTGAGGCTCTCCTGTAAAGCCTCTGGACGTACCGTCAACCAGAACATGGCGTGGTTCCGCCAGCCTCCAGGGAAGGAGCGTGAGTTTGTGTCAGCTATTGAGTGGAGTGTTGGAATGACAAGATATAAAGACTCCGTGAAGGGCCGATTCACCATCTCCAGAGACATCGCCAAGGGCACGGTGTATCTGCAAATGAACAGCCTGAAACCTGACGACACGGCCGTTTATTACTGTGCAGCAAGTAACTCAGGCGTTGGCGCGTATGGGTATGATTATTGGGGCCAGGGGACCCAGGTCACCGTCTCCTCA |
| 6 | BA.1_F2 | CAGGTGCAGCTGCAGGAGTCTGGGGGAGGATTGGTGCAGGCAGGGGGCTCTCTGAGACTCTCCTGTGCAGCCTCTGGACTGCGATTCAGTAACTATAACATGGGCTGGTTCCGCCGGGCTCCAGGGAAGGACCGTGAGTTTGTAGCATATATTAGCTGGAGTGATGATACCACAGCTTATGCAGACTCCGTGAAGGGCCGATTCACCATCTCCAGAGACAACGCCAAGAACACGGTGTATCTGCAAATGAACAGCCTGAATTCTGAGGACACGGCCGTTTACTACTGTGCAGCAGTTGGTGGTTACTACCAGGCGGGAGATCGGCCCTCGTCGGAATATGAGTATGACTACTGGGGTCAGGGGACCCGGGTCACCGTCTCCTCA |

| | | |
|---|---|---|
| 7 | BA.1_A1 | CAGGTGCAGCTGCAGGAGTCTGGGGGAGGCTTGGTGCAGCCTGGGGGT<br>TCTCTGAGACTCTCCTGTGCGACCTCTGGATTCACTTTGGATAATTATGCC<br>ATAGGCTGGTTCCGCCAGGCCCCAGGGAAGGAGCGCGAGGGGGTCTCAT<br>GTATTAGGAGTAGTGATGGTACCACATACTATGCAGATTCCGTGAAGGGC<br>CGATTCACCATGTCCAGTGACAACGCCAAGAATATGTATCTGCAAATGAA<br>TAACCTGAAACCCGAGGACACGGCCGTTTATTACTGTGCAGCAGGGGGT<br>CAAACGTGTTCGGAGACGGTAGTACGTGGTTGGCTGCTGGGTGACTACT<br>GGGGCCAGGGGACCCAGGTCACCGTCTCCTCA |
| 8 | BA.1_C2 | CAGGTGCAGCTGCAGGAGTCTGGGGGAGGCTTGGTGCAGACTGGGGAC<br>TCTCTGAGACTCTCCTGTGTAGCCTCTGGACTGTACTTCAGGTACCATGC<br>CATGGGCTGGTTCCGCCAGGCTCCAGGAAAGGAGCGTGAATTTATAGCA<br>GGTATTAGCGGGAATGGTAGAACCACGGACTATGCAGACTCCGTGAACG<br>GCCGATTCACCATCTCCAGAGACAACGACAAGAACACGATGTACCTGCA<br>AATGAACAGCCTGAAACCTGACGACACGGCCGTTTATTACTGCGCTGGC<br>CGTGGTCGCAATTTTGTTATAATCGCGCGACACAGGGCGGAGTATGTCCA<br>CTGGGGCCAGGGGACCCAGGTCACCGTCTCCTCA |
| 9 | BA.1 _E11 | CAGGTGCAGCTGCAGGAGTCTGGGGGAGGATTGGTGCAGGCTGGGGAC<br>TCTCTGAACCTCTCCTGTGTAGTGTCTGGAGGTACTTTCAGCAGGTATAC<br>CCTGGGCTGGTTCCGCCAGGCTCCAGGGAAGGAGCGCGAGTTTGTAAGT<br>GGTATTAATTGGAGTGGTATCAGCGCAACAATTCCAGCCGTGAAGGGCCG<br>GTTCACCATCGGGAGAGACAACACCAAGAACACGGGATATTTGCAAATG<br>CACAGGTTGGAACCTGAGGACACGGCCGTTTATTACTGTGCATTAGACAC<br>GACATTTCCATCTGGCGCCTTGACTGAAGCTTCAGAATATGACTACTGGG<br>GCCAGGGGACCCAGGTCACCGTCTCCTCA |
| 10 | BA.1_D3 | CAGGTGCAGCTGCAGGAGTCTGGGGGAGGCTCGGTGCAGCCTGGGGGA<br>TCTCTGAGACTCTCCTGTGCAGCCTCTGGGTTCACCTTCGGTATTTATGGC<br>ATGACCTGGGTCCGTCAGGCTCCAGGAAAGGGGCACGAATGGATCTCTA<br>CTATTACCGCCGGTGGTGAGATTACCCACTATGCAGACTCCGTTCGGGGC<br>CGATTCTCCATCTCCAGAGACAACGCCAAAAATACGCTGTATTTGGAGAT<br>GAACAGCCTGAAACTGGAGGACACGGCCCGTTATTACTGTGCACGGACT<br>CCTGGCATAGTAGTACGTGGACCCAATACATACGACTACCTGGGCCAGGG<br>AACCCAGGTCACCGTCTCCTCA |
| 11 | BA.1_D10 | CAGGTGCAGCTGCAGGAGTCTGGGGGAGGCTCGGTGCAGGCTGGGGGG<br>TCTCTGAGACTCTCCTGTGCAGCCTCGGGATTCACCTTCGGTATTTATGGT<br>ATGAGCTGGCTCCGCCAGGCTCCAGGAAAGGGGCGAGAGTGGGTCTCA<br>ACTATTACCGCTGGCGGTGAGATCCAACACTATGCAGACTCCGTGAAGG<br>GCCGATTCACCATCTCCAAAGACAACGCCAAGAACATGCTGTATCTGCA<br>AATGACTAGCCTGGAAGTTGAGGACACGGCCGTTTATTACTGTGCACGA<br>GTTCCAGGGATAGTAGTACGGGGGTCCAATGCGTATGACCACGTGGGTC<br>AGGGGACCCAGGTCACCGTCTCCTCA |
| 12 | BA.1 _E7 | CAGGTGCAGCTGCAGGAGTCTGGAGGACGACTGGCGCAGCCTGGGGAC<br>TCTCTGAGACTCTCCTGTGCAGCCTCTGGCCTCAACTTCAGCGATTACAC<br>CATGGGATGGTTCCGCCAGGCTCCAGGAGAGGAGCGTGAATTAGTGGCG<br>CATATCACCCTGACTGGTTTTACATTGCGTGGTAGAAGTCCATACTATGCA<br>GACTCCGTGAAGGACCGATTCACCATCTCCAGAGACATTGTCAAGAACG<br>CGGTTTATCTGCAAATGAACAACCTGAAATTTGAGGATTCGGCTGTTTAT<br>TACTGTGCAGCAGGTGCTTACCCACTCTCATTGACGGCTGTTGCCTACTG<br>GGGCCAGGGGACCCAGGTCACCGTCTCCTCA |

| | | |
|---|---|---|
| 13 | BA.1_G9 | CAGGTGCAGCTGCAGGAGTCTGGGGGAGGATTGGTGCAGCCTGGGGGCTCTCTGAGACTCTCCTGTGCAATCTCCGGACGCGCCTCCGATTTCTATGCCATGGGCTGGTTCCGCCAAACTCCAGGAGAGGAACGTGAGTTTCTAGCAGCCATTACCTTGAAGACTTTTCGCACGCGCTATGCGGCCTCCGTGGAGGGTCGATTCCGCTTCTCCAGAGACAATCCCGAGAACACGGTATATCTGCAATTGAACGAACTGACACCTGACGATACGGCCGTTTATTACTGTGGTCTTACCGAGGTTATGAGTCTTTCGCCAATGCCACATGACTATCAGTACTGGGGCCAGGGGACCCAGGTCACCGTCTCCTCA |
| 14 | MERS_C9 | CAGGTGCAGCTGCAGGAGTCTGGGGGAGGATTGGTCCAGGCTGGGGGCTCTCTGAGACTCTCCTGTGCAGCCACTGGACGCGGTTTCAGTGACAGAGCCATGGGCTGGTTCCGCCAGGCTCCAGGGAAGGGGCGTGAGTTTGTGGCTGCTATTAACATGGGTGCCTTTGACACAGTGTATGGAGACTCTGTCAAGGACCGATTCGCCATCTCCAGAGACGACGCCAAGAACACAATGTATCTGCAAATGAACAGCCTGATACCTGAGGACACGGCCGTTTATTACTGTGCAGTCGGGTATGGATCGTTCCTCAGTCGTAATCAATATTCTTATGAAGTCTGGGGCCAGGGGACCCAGGTCACCGTCTCCTCA |
| 15 | BA.1_B3 | CAGGTGCAGCTGCAGGAGTCTGGGGGAGGATTGGTGCAGGCTGGGGGCTCTCTGAGACTCTCCTGTGGAGTCTCTGGAGGCTACTTCACTAACTATGCCATGGGCTGGTTCCGCCAGCCTCCCGGGAAGGAGCGGAGTGAATTTGTGGCAGGCATTCAGTGGAGTGGCGGTTACGAATATTATTTCGACTCCGTGAAGGGCCGATTCGCCATCTCCACAGACAACGCCAGGAACACGGTGTATCTGCAAATGAACAGCCTGAAACCTGAGGACACGGCCGTTTATTACTGCGCAGCCGCGAAGCTCGAACATTACGATAGCCACTTTCCAATGGAGTCATATGAGTATAACTACTGGGGCCAGGGGACCCAGGTCACCGTCTCCTCA |
| 16 | BA.1 _E1 | CAGGTGCAGCTGCAGGAGTCTGGGGGAGGATTGGTGCAGGCTGGGGACTCTCTGACACTGTCCTGTGCGGTCTCTGGACGCACCTCCGGCACCACCATGGCCTGGTTCCGCCAGGCTCCAGGGAAGGATCGTGACTTTGTAGGCGCTATTAATTGGTATGTTGGCGGCCCTCACTATGCAGACTCCGTTAAGGGCCGATTCAGTATCTCGAGAGACAACACCAACAACATCTTGTATCTGCAAATGAACACCCTACAGCCTGACGACACGGCCGTTTATTACTGTGCAGCAAAAGACTGGCAACCGGCCCTCAAATCACGGCCAGATGACTATCCCTACTGGGGCCAGGGGACCCAGGTCACCGTCTCCTCA |
| 17 | BA.1 _C1 | CAGGTGCAGCTGCAGGAGTCTGGGGGAGGATTGGTGAAGGCTGGGGACTCTCTGACACTCTCCTGTGCAGTCTCTGGACGCACCTCCGGCACCACCATGGCCTGGTTCCGCCAGGCGCCAGAGAAGGACCGTGAGTTTGTAGCTGCTATTAACTGGAACTTTGGTGGCCCACACTATGCAGACTCCGTGCAGGGCCGATTCACCATCTCCAGAAACAACGCCGAGAATACACTGACTCTGCAAATGAACATCCTGGAACCTGACGACACGGCCGTTTATTACTGTGCAGCAAGAGACTGGACACCGGGCCTCAAAACGCGACCAGATGACTATGCCTACTGGGGCCAGGGGATTCAGGTCACCGTCTCCTCG |
| 18 | BA.1_B1 | CAGGTGCAGCTGCAGGAGTCTGGGGGAGGATTGGTGCAGACTGGGGGCTCCCTGAGACTCTCCTGTGCAGCCTCTGGACGCACCTCTAGTAGGTATGACATGGACTGGTACCGCCAGGCTCCAGGGAAGGAGCGTGAGTTCGTAGCGGGTTTCAGCCGGAATGGTATTAGCACATATTATGAAGACTCCGTGAAGGGCCGATTCACCATCTCCAGAGACAACGCCAAGAACACAGTGTATCTGCAAATGAGCAGCCTGAAACCTGAGGACACGGCCGTTTATTACTGTGGAGCACGTGTACGTGGCAGCTTAACGTATGATTCGTGGGGCCAGGGGACCCAGGTCACCGTCTCCTCA |

| | | |
|---|---|---|
| 19 | MERS_B8 | CAGGTGCAGCTGCAGGAGTCTGGGGGAGGATTGGTGCAGGCTGGGGGC<br>TCTCTGAGACTCTCCTGTGCAGCCTCCGGACGCACCTTTAGTAGCTATAC<br>AATGGGGTGGTTCCGCCAGGCTCCAGGGAAGGAGCGTGAGTTTGTAGCA<br>GCTATTAATAGGAGTGGAGATAGGACATCGTACGCAGACTCCGCGAAGG<br>GCCGATTTACTATCTCCAGAGACAACGCCAAGAACACGGTGTATCTGCAA<br>ATGAACAGCCTGAAACCTGAGGACACGGCCACTTATTACTGTGCAGCGC<br>ACGAATCTGCTAATGCTCAGGCTATGGCTGTTATGAGGGGTCGTGGGATT<br>AACTACTGGGGCCAGGGGACCCAGGTCACCGTCTCCTCA |
| 20 | MERS_F6 | CAGGTGCAGCTGCAGGAGTCTGGGGGAGGATTGGTGCAGGCTGGGGAC<br>TCTCTAGGACTCTCCTGTGTAGCCTCCGGACGCGGCCTAGATTCTCTAAT<br>AGTGGCCTGGTTCCGCCAGGCTCCCGGAAAGGAGCGTGAGTTTGTAGCA<br>GGTATCGTCTGGAGTGATGAATTTACATCGTATGGCAAGTTCGCGCAGGG<br>CCGATTTACCATCTCCAGAGACAAGGGCAAGAACACGATATTTCTGCAA<br>ATTAACAGCCTGAAACCTGAGGACACGGCCGTTTATTACTGTGCCGGAC<br>GTTACGGGAATCTTATTCATGAAAACGAGAATGAGTACCAAGTTTGGGGC<br>CAGGGGACCCAGGTCACCGTCTCCTCA |
| 21 | MERS_D1 | CAGGTGCAGCTGCAGGAGTCTGGGGGAGGATTGGTGCAGGCTGGGGAC<br>TCTCTGAGACTCACCTGTACAGCCTCTGGATATATTCACGAGACGCATGT<br>CGTGGGCTGGTTCCGCCAGGCTCCAGGAAAGGAGCGTGAGTTTGTGGC<br>ACATATTACATGGGGTCTTGGCTACACAGCCTATGAGGACGCCGTGAAGG<br>GCCGCTTCACCATTACCAGAGACAACGCCAAAAACACGATTTATCTGCA<br>AATGAACAGCCTGAAACCTGAGGACACGGCCAGATATTACTGTGCAGTT<br>CGGCCAGGCGGGATTCACTTTGGTTCCTGGGGTCCGGGGACCCAGGTCA<br>CCGTCTCCTCA |
| 22 | MERS_G9 | CAGGTGCAGCTGCAGGAGTCTGGGGGAGGATTGGTACAGGCTGGGGGC<br>TCTTTGACACTCTCCTGTGCAGCCTCTGGACTACCCTTCAGTACATATACC<br>GTGGGCTGGTTCCGCCAGGCTCCAGGGAAGGAACGTGAATTTGTAGCGC<br>GGATTACTCGGAACATTTATAACACAATTTATGCAGATTCCGTACAGGGCC<br>GCTTCACCATCTCCAGAGACACCACCAAAAACACGGTGTATCTGCAAAT<br>GAACAGCCTGAAATTTGAGGACACGGCCGTTTATTTCTGCGCAGCGCGC<br>CCGTCCGGAAGTACCATGATAGCCTCAGACTATGACTACTGGGGCCAGG<br>GGACCCAGGTCACCGTCTCCTCA |
| 23 | MERS_A8 | CAGGTGCAGCTGCAGGAGTCTGGGGGAGAATTCGTGCAGCCCGGGGAC<br>TCGCTGAGACTCTCCTGCGCGACCTCTGGATTCACTTTGGATAATTATGCC<br>ATAGGCTGGTTCCGCCAGGCCCCAGGGAAGGAGCGCGAGGGGGTCTCAT<br>GTATTAGGAGTAGTGATAGTACCACATACTATGCAGATTCCGTGAAGGGC<br>CGATTCACCATGTCCAGTGACAACGCCAAGAATATGTATCTGCAAATGAA<br>CAACCTGAAACCCGAGGACACGGCCGTTTATTACTGTGCAGCAGGGGGT<br>CAAACGTGTTCGGAGACGGTAGTACGTGGTTGGCTGCTGGGTGACTACT<br>GGGGCCAGGGGACCCAGGTCACCGTCTCCTCG |
| 24 | MERS_A4 | CAGGTGCAGCTGCAGGAGTCTGGGGGAGAATTCGTGCAGCCCGGGGAC<br>TCGCTGAGACTCTCCTGCGCGACCTCTGGATTCACTTTGGATAATTATGCC<br>ATAGGCTGGTTCCGCCAGGCCCCAGGGAAGGAGCGCGAGGGGGTCTCAT<br>GTATTAGGAGTAGTGATAGTACCACATACTATGCAGATTCCGTGAAGGGC<br>CGATTCACCATGTCCAGTGACAACGCCAAGAATATGTATCTGCAAATGAA<br>CAACCTGAAACCCGAGGACACGGCCGTTTATTACTGTGCAGCAGGGGGT<br>CAAACGTGTTCGGAGACGGTAGTACGTGGTTGGCTGCTGGGTGACTACT<br>GGGGCCAGGGGACCCAGGTCACCGTCTCCTCG |

**Table 2** VHH amino acid sequences (nos. 1-13 from BA.1 S2 panning only; nos. 14-18 from BA.1 S2 and MERS S2 panning and nos. 19-24 from MERS S2 panning only)

| Number | VHH id | CDR1-IMGT | CDR2-IMGT | CDR3-IMGT | Amino acid sequence |
|---|---|---|---|---|---|
| 1 | BA.1_D12 | GRGFDAYG | INWGGTT | AAAPTSTVVTRPSRPKY | QVQLQESGGGLVQPGDSLTLSCAASGRGFDAYGMMWFRQAPGKEREFVTGINWGGTTYYADSAKGRFTISRTSDKTTVYLQMTSLKPEDSAIYYCAAAPTSTVVTRPSRPKYWGQGTQVTVSS |
| 2 | BA.1 _A7 | GRTVSLFD | ITLKEGKL | AADQTVVRMTGTGIDY | QVQLQESGGGSVQAGGSLRLSCVASGRTVSLFDMGWFRQAPGKEREFVARITLKEGKLNYADSVKGRFTISRGNPENTVYLQMDSLKPEDTAVYYCAADQTVVRMTGTGIDYWGQGTQVTVSS |
| 3 | BA.1 _D2 | GRAFSVTT | SDARGGK | AAGPFGVSAREDDYAY | QVQLQESGGGVVQAGGSLRLSCAASGRAFSVTTVAWFRQSPGKEREYIARSDARGGKYYTDSVKGRFTISDERDKMTVYLQMNDLETSDTAVYYCAAGPFGVSAREDDYAYWGQGTQVTVSS |
| 4 | BA.1 _A5 | GRILSNYV | ITWSAGRT | AARIVSASEYTY | QLQESGGGLVQAGGSLRLSCATSGRILSNYVMGWFRQAPGKERALLGAITWSAGRTAYAESVKGRFTISRDIAENAVYLQMNSLKPEDTAVYYCAARIVSASEYTYWGQGTQVTVSS |
| 5 | BA.1 _B10 | GRTVNQNM | EWSVGMT | AASNSGVGAYGYDY | QVQLQESGGGLVQAGGSLRLSCKASGRTVNQNMAWFRQPPGKEREFVSAIEWSVGMTRYKDSVKGRFTISRDIAKGTVYLQMNSLKPDDTAVYYCAASNSGVGAYGYDYWGQGTQVTVSS |
| 6 | BA.1 _F2 | GLRFSNYN | ISWSDDTT | AAVGGYYQAGDRPSSEYEYDY | QVQLQESGGGLVQAGGSLRLSCAASGLRFSNYNMGWFRRAPGKDREFVAYISWSDDTTAYADSVKGRFTISRDNAKNTVYLQMNSLNSEDTAVYYCAAVGGYYQAGDRPSSEYEYDYWGQGTRVTVSS |

| | | | | | |
|---|---|---|---|---|---|
| 7 | BA.1 _A1 | GFTLDNYA | IRSSDGTT | AGGQTCSETVVRGWLLGDY | QVQLQESGGGLVQPGGSLRLSCATSGFTLDNYAIGWFRQAPGKEREGVSCIRSSDGTTYYADSVKGRFTMSSDNAKNMYLQMNNLKPEDTAVYYCAAGGQTCSETVVRGWLLGDYWGQGTQVTVSS |
| 8 | BA.1_C2 | GLYFRYHA | ISGNGRTT | AGRGRNFVIIARHRAEYVH | QVQLQESGGGLVQTGDSLRLSCVASGLYFRYHAMGWFRQAPGKEREFIAGISGNGRTTDYADSVNGRFTISRDNDKNTMYLQMNSLKPDDTAVYYCAGRGRNFVIIARHRAEYVHWGQGTQVTVSS |
| 9 | BA.1_E11 | GGTFSRYT | INWSGIS | ALDTTFPSGALTEASEYDY | QVQLQESGGGLVQAGDSLNLSCVVSGGTFSRYTLGWFRQAPGKEREFVSGINWSGISATIPAVKGRFTIGRDNTKNTGYLQMHRLEPEDTAVYYCALDTTFPSGALTEASEYDYWGQGTQVTVSS |
| 10 | BA.1_D3 | GFTFGIYG | ITAGGEIT | ARTPGIVVRGPNTYDY | QVQLQESGGGSVQPGGSLRLSCAASGFTFGIYGMTWVRQAPGKGHEWISTITAGGEITHYADSVRGRFSISRDNAKNTLYLEMNSLKLEDTARYYCARTPGIVVRGPNTYDYLGQGTQVTVSS |
| 11 | BA.1_D10 | GFTFGIYG | ITAGGEIQ | ARVPGIVVRGSNAYDH | QVQLQESGGGSVQAGGSLRLSCAASGFTFGIYGMSWLRQAPGKGREWVSTITAGGEIQHYADSVKGRFTISKDNAKNMLYLQMTSLEVEDTAVYYCARVPGIVVRGSNAYDHVGQGTQVTVSS |
| 12 | BA.1_E7 | GLNFSDYT | ITLTGFTL | AAGAYPLSLTAVAY | QVQLQESGGRLAQPGDSLRLSCAASGLNFSDYTMGWFRQAPGEERELVAHITLTGFTLRGRSPYYADSVKDRFTISRDIVKNAVYLQMNNLKFEDSAVYYCAAGAYPLSLTAVAYWGQGTQVTVSS |
| 13 | BA.1_G9 | GRASDFYA | ITLKTFRT | GLTEVMSLSPMPHDYQY | QVQLQESGGGLVQPGGSLRLSCAISGRASDFYAMGWFRQTPGEEREFLAAITLKTFRTRYAASVEGRFRFSRDNPENTVYLQLNELTPDDTAVYYCGLTEVMSLSPMPHDYQYWGQGTQVTVSS |

| | | | | | |
|---|---|---|---|---|---|
| 14 | MERS_C9 | GRGFSDRA | INMGAFDT | AVGYGSFLSRNQYSYEV | QVQLQESGGGLVQAGGSLRLSCAATGRGFSDRAMGWFRQAPGKGREFVAAINMGAFDTVYGDSVKDRFAISRDDAKNTMYLQMNSLIPEDTAVYYCAVGYGSFLSRNQYSYEVWGQGTQVTVSS |
| 15 | BA.1_B3 | GYFTNYAM | IQWSGGYE | AAAKLEHYDSHFPMESYEYNY | QVQLQESGGGLVQAGGSLRLSCGVSGGYFTNYAMGWFRQPPGKERSEFVAGIQWSGGYEYYFDSVKGRFAISTDNARNTVYLQMNSLKPEDTAVYYCAAAKLEHYDSHFPMESYEYNYWGQGTQVTVSS |
| 16 | BA.1_E1 | GRTSGTTM | NWYVGGP | AAKDWQPALKSRPDDYPY | QVQLQESGGGLVQAGDSLTLSCAVSGRTSGTTMAWFRQAPGKDRDFVGAINWYVGGPHYADSVKGRFSISRDNTNNILYLQMNTLQPDDTAVYYCAAKDWQPALKSRPDDYPYWGQGTQVTVSS |
| 17 | BA.1_C1 | GRTSGTTM | NWNFGGP | AARDWTPGLKTRPDDYAY | QVQLQESGGGLVKAGDSLTLSCAVSGRTSGTTMAWFRQAPEKDREFVAAINWNFGGPHYADSVQGRFTISRNNAENTLTLQMNILEPDDTAVYYCAARDWTPGLKTRPDDYAYWGQGIQVTVSS |
| 18 | BA.1_B1 | GRTSSRYD | FSRNGIST | GARVRGSLTYDS | QVQLQESGGGLVQTGGSLRLSCAASGRTSSRYDMDWYRQAPGKEREFVAGFSRNGISTYYEDSVKGRFTISRDNAKNTVYLQMSSLKPEDTAVYYCGARVRGSLTYDSWGQGTQVTVSS |
| 19 | MERS_B8 | GRTFSSYT | INRSGDRT | AAHESANAQAMAVMRGRGINY | QVQLQESGGGLVQAGGSLRLSCAASGRTFSSYTMGWFRQAPGKEREFVAAINRSGDRTSYADSAKGRFTISRDNAKNTVYLQMNSLKPEDTATYYCAAHESANAQAMAVMRGRGINYWGQGTQVTVSS |

| | | | | | |
|---|---|---|---|---|---|
| 20 | MERS_F6 | GRGLDSLI | IVWSDEFT | AGRYGNLIHENENEYQV | QVQLQESGGGLVQAGDSLGLSCVASGRGLDSLIVAWFRQAPGKEREFVAGIVWSDEFTSYGKFAQGRFTISRDKGKNTIFLQINSLKPEDTAVYYCAGRYGNLIHENENEYQVWGQGTQVTVSS |
| 21 | MERS_D1 | GYIHETHV | ITWGLGYT | AVRPGGIHFGS | QVQLQESGGGLVQAGDSLRLTCTASGYIHETHVVGWFRQAPGKEREFVAHITWGLGYTAYEDAVKGRFTITRDNAKNTIYLQMNSLKPEDTARYYCAVRPGGIHFGSWGPGTQVTVSS |
| 22 | MERS_G9 | GLPFSTYT | ITRNIYNT | AARPSGSTMIASDYDY | QVQLQESGGGLVQAGGSLTLSCAASGLPFSTYTVGWFRQAPGKEREFVARITRNIYNTIYADSVQGRFTISRDTTKNTVYLQMNSLKFEDTAVYFCAARPSGSTMIASDYDYWGQGTQVTVSS |
| 23 | MERS_A8 | GFTLDNYA | IRSSDSTT | AGGQTCSETVVRGWLLGDY | QVQLQESGGEFVQPGDSLRLSCATSGFTLDNYAIGWFRQAPGKEREGVSCIRSSDSTTYYADSVKGRFTMSSDNAKNMYLQMNNLKPEDTAVYYCAAGGQTCSETVVRGWLLGDYWGQGTQVTVSS |
| 24 | MERS_A4 | GFTLDNYA | IRSSDSTT | AGGQTCSETVVRGWLLGDY | QVQLQESGGEFVQPGDSLRLSCATSGFTLDNYAIGWFRQAPGKEREGVSCIRSSDSTTYYADSVKGRFTMSSDNAKNMYLQMNNLKPEDTAVYYCAAGGQTCSETVVRGWLLGDYWGQGTQVTVSS |

**Supplementary Table 2.** Sequences of beta-coronavirus spike and S2 proteins

| Protein | DNA sequence | Amino acid sequence |
|---|---|---|
| SARS-CoV-2<br>BA.1 S (6 pro) | ATGTTCGTGTTCCTGGTGCTCCTGCCTCTGGTGAGCAGCCAGTGCGTG<br>AACCTGACCACCCGAACCCAGCTCCCACCAGCCTACACCAACAGCTTT<br>ACACGGGGCGTGTACTACCCTGACAAGGTGTTCAGATCTAGCGTCCTG<br>CACAGCACTCAGGACCTCTTCCTGCCGTTCTTCAGCAACGTGACATGG<br>TTCCACGTGATCAGCGGCACAAACGGAACCAAGCGGTTTGATAACCCC<br>GTCCTGCCATTCAATGATGGAGTTTACTTCGCCAGTATCGAGAAGAGT<br>AACATCATCCGGGGCTGGATCTTCGGCACCACCCTGGATAGCAAAACA<br>CAGAGCCTCCTGATCGTGAACAATGCCACGAACGTCGTGATCAAGGTG<br>TGCGAGTTCCAGTTTTGCAATGATCCTTTCCTGGATCACAAGAACAAC<br>AAGAGCTGGATGGAAAGCGAGTTCAGAGTCTACAGCAGCGCCAACAAC<br>TGCACATTCGAGTACGTCTCTCAGCCTTTTCTGATGGACCTTGAGGGG<br>AAACAAGGCAACTTCAAGAACCTGAGAGAATTCGTGTTCAAGAACATC<br>GACGGCTACTTCAAAATCTACTCCAAGCACACACCCATCATAGTCCGG<br>GAACCTGAGGACCTCCCTCAGGGCTTCAGCGCCCTGGAACCCCTGGTC<br>GACCTGCCCATAGGCATCAACATAACGCGGTTCCAAACCCTGCTGGCC<br>CTGCATAGATCCTACCTGACTCCTGGCGACAGCAGCAGCGGATGGACC<br>GCCGGAGCTGCAGCCTACTATGTGGGCTACCTGCAACCTAGAACCTTC<br>CTGCTGAAGTACAACGAGAACGGCACAATCACAGACGCCGTCGACTGC<br>GCCCTGGACCCTCTCTCTGAGACAAAGTGCACCCTGAAGTCCTTCACC<br>GTGGAAAAGGGCATCTACCAGACCAGCAACTTCCGGGTGCAGCCTACA<br>GAGAGCATCGTGCGATTTCCAAACATTACCAACCTCTGCCCCTTCGAC<br>GAGGTGTTTAACGCCACAAGATTTGCCTCCGTTTACGCCTGGAATAGA<br>AAGAGAATCAGCAATTGTGTGGCCGACTACTCCGTGCTGTATAACCTG<br>GCCCCATTCTTCACCTTCAAGTGCTACGGCGTTTCCCCAACAAAGCTG<br>AATGACCTGTGCTTCACCAACGTGTACGCCGACTCCTTCGTAATTAGA<br>GGCGATGAGGTGCGGCAGATCGCACCAGGCCAGACCGGTAACATCGCT<br>GACTACAACTATAAGCTGCCTGATGATTTTACAGGCTGCGTGATCGCC<br>TGGAACTCTAACAAGCTGGATAGCAAGGTGTCTGGCAACTACAACTAC<br>CTGTACCGGCTGTTTCGCAAGTCTAACCTGAAACCTTTCGAGAGAGAC<br>ATCTCCACAGAGATCTACCAGGCCGGTAACAAGCCTTGTAACGGGGTG<br>GCCGGCTTCAACTGTTACTTCCCTCTGAAGAGCTACTCCTTCCGGCCT<br>ACCTACGGAGTCGGCCACCAGCCATACCGGGTGGTCGTGCTGTCCTTC<br>GAGTTACTCCACGCCCCCGCCACCGTCTGCGGTCCTAAGAAGTCCACC<br>AATCTGGTTAAGAACAAATGCGTGAACTTCAACTTCAACGGCCTGAAG<br>GGGACCGGCGTGCTGACCGAAAGCAACAAAAAGTTCCTCCCCTTCCAG<br>CAGTTCGGCCGTGATATCGCTGACACCACAGACGCCGTCAGAGATCCA<br>CAGACCCTGGAAATCCTGGATATTACACCCTGCTCCTTCGGAGGAGTT<br>TCTGTGATCACCCCCGGGACCAATACCAGCAACCAGGTGGCTGTGCTG<br>TACCAAGGCGTTAACTGCACCGAGGTTCCTGTGGCCATCCACGCCGAT<br>CAGCTGACACCTACTTGGAGAGTGTACTCCACTGGCTCCAATGTGTTC<br>CAGACCAGGGCCGGATGTCTGATCGGCGCCGAGTACGTGAATAACAGT<br>TACGAGTGCGACATCCCTATCGGCGCCGGCATCTGTGCCAGCTACCAG<br>ACCCAGACAAAGAGCCACGGGTCTGCTTCCTCTGTAGCTAGCCAGAGC<br>ATCATCGCCTACACCATGAGCCTGGGCGCCGAGAACAGCGTGGCCTAT<br>TCCAACAACTCTATCGCCATTCCCACCAACTTTACAATTAGCGTCACA<br>ACAGAGATCCTGCCCGTGAGCATGACCAAGACCAGCGTGGACTGTACA<br>ATGTACATCTGTGGCGACAGCACTGAATGCAGCAACCTGCTGCTGCAA<br>TACGGCTCCTTTTGCACCCAACTGAAGCGGGCGCTGACCGGAATCGCC<br>GTGGAACAGGACAAAAATACCCAGGAGGTGTTCGCCCAAGTGAAGCAG<br>ATCTACAAGACCCCACCTATCAAGTACTTCGGCGGCTTTAACTTTAGC<br>CAGATTCTCCCTGATCCTTCTAAGCCTAGCAAGCGGAGCCCTATCGAG<br>GATCTGCTGTTCAACAAGGTCACCCTGGCCGATGCCGGCTTTATCAAA<br>CAGTATGGCGATTGCCTGGGCGACATAGCCGCCAGAGATCTGATCTGC<br>GCCCAGAAATTCAAGGGCCTGACAGTTCTCCCACCTCTGCTGACCGAC | MFVFLVLLPLVSSQCVNLTTRTQLPPAYT<br>NSFTRGVYYPDKVFRSSVLHSTQDLFLPF<br>FSNVTWFHVISGTNGTKRFDNPVLPFNDG<br>VYFASIEKSNIIRGWIFGTTLDSKTQSLL<br>IVNNATNVVIKVCEFQFCNDPFLDHKNNK<br>SWMESEFRVYSSANNCTFEYVSQPFLMDL<br>EGKQGNFKNLREFVFKNIDGYFKIYSKHT<br>PIIVREPEDLPQGFSALEPLVDLPIGINI<br>TRFQTLLALHRSYLTPGDSSSGWTAGAAA<br>YYVGYLQPRTFLLKYNENGTITDAVDCAL<br>DPLSETKCTLKSFTVEKGIYQTSNFRVQP<br>TESIVRFPNITNLCPFDEVFNATRFASVY<br>AWNRKRISNCVADYSVLYNLAPFFTFKCY<br>GVSPTKLNDLCFTNVYADSFVIRGDEVRQ<br>IAPGQTGNIADYNYKLPDDFTGCVIAWNS<br>NKLDSKVSGNYNYLYRLFRKSNLKPFERD<br>ISTEIYQAGNKPCNGVAGFNCYFPLKSYS<br>FRPTYGVGHQPYRVVVLSFELLHAPATVC<br>GPKKSTNLVKNKCVNFNFNGLKGTGVLTE<br>SNKKFLPFQQFGRDIADTTDAVRDPQTLE<br>ILDITPCSFGGVSVITPGTNTSNQVAVLY<br>QGVNCTEVPVAIHADQLTPTWRVYSTGSN<br>VFQTRAGCLIGAEYVNNSYECDIPIGAGI<br>CASYQTQTKSHGSASSVASQSIIAYTMSL<br>GAENSVAYSNNSIAIPTNFTISVTTEILP<br>VSMTKTSVDCTMYICGDSTECSNLLLQYG<br>SFCTQLKRALTGIAVEQDKNTQEVFAQVK<br>QIYKTPPIKYFGGFNFSQILPDPSKPSKR<br>SPIEDLLFNKVTLADAGFIKQYGDCLGDI<br>AARDLICAQKFKGLTVLPPLLTDEMIAQY<br>TSALLAGTITSGWTFGAGPALQIPFPMQM<br>AYRFNGIGVTQNVLYENQKLIANQFNSAI<br>GKIQDSLSSTPSALGKLQDVVNHNAQALN<br>TLVKQLSSKFGAISSVLNDIFSRLDPPEA<br>EVQIDRLITGRLQSLQTYVTQQLIRAAEI<br>RASANLAATKMSECVLGQSKRVDFCGKGY<br>HLMSFPQSAPHGVVFLHVTYVPAQEKNFT<br>TAPAICHDGKAHFPREGVFVSNGTHWFVT<br>QRNFYEPQIITTDNTFVSGNCDVVIGIVN<br>NTVYDPLQPELDSFKEELDKYFKNHTSPD<br>VDLGDISGINASVVNIQKEIDRLNEVAKN<br>LNESLIDLQELGKYEQGSGYIPEAPRDGQ<br>AYVRKDGEWVLLSTFLGRS |

| | | |
|---|---|---|
| | GAGATGATCGCTCAGTACACCTCTGCCCTGCTGGCTGGCACCATCACA<br>TCTGGGTGGACATTTGGCGCCGGCCCCGCCCTGCAGATCCCCTTTCCC<br>ATGCAGATGGCCTATAGATTCAACGGAATCGGCGTGACCCAGAACGTG<br>CTGTATGAAAACCAGAAGCTGATCGCTAACCAGTTCAATTCTGCCATC<br>GGCAAGATCCAGGACTCCCTCTCCTCTACCCCCAGCGCCCTGGGCAAA<br>CTGCAGGACGTGGTGAATCACAACGCCCAAGCCCTGAACACCCTGGTG<br>AAGCAGCTCAGCAGCAAGTTTGGCGCCATCAGCTCTGTGCTGAACGAT<br>ATCTTCTCTAGACTGGACCCTCCAGAAGCCGAAGTCCAGATCGATAGA<br>CTGATCACAGGCAGACTGCAGTCCCTGCAAACCTACGTGACCCAACAG<br>CTGATCAGGGCCGCTGAAATAAGAGCCAGCGCCAATCTCGCCGCTACC<br>AAGATGTCCGAGTGTGTGCTGGGACAGTCTAAACGCGTTGACTTCTGC<br>GGCAAAGGCTATCACCTGATGAGCTTCCCCCAGAGCGCGCCGCACGGC<br>GTGGTGTTCCTGCATGTGACATACGTGCCTGCCCAAGAGAAGAATTTC<br>ACAACCGCCCCTGCCATCTGCCACGACGGCAAGGCCCACTTCCCAAGA<br>GAGGGCGTTTTCGTTTCCAATGGCACACACTGGTTCGTGACACAAAGA<br>AACTTCTACGAACCCCAGATTATCACCACCGACAACACCTTCGTGAGT<br>GGCAATTGTGACGTGGTCATCGGAATCGTGAACAACACAGTGTACGAC<br>CCTCTGCAACCTGAGCTGGACTCTTTTAAGGAAGAGCTGGACAAGTAC<br>TTTAAAAACCACACCAGCCCCGATGTGGACCTGGGCGACATCAGTGGC<br>ATTAACGCCAGCGTGGTGAACATCCAAAAGGAAATCGACAGACTGAAC<br>GAGGTGGCCAAGAACCTGAACGAGTCCCTGATCGACCTGCAGGAGCTC<br>GGCAAATACGAGCAGGGATCCGGATACATCCCCGAGGCCCCCAGAGAT<br>GGCCAGGCCTACGTGCGGAAGGACGGCGAGTGGGTACTGCTGAGCACA<br>TTCCTGGGCAGATCC | |
| SARS-CoV-2 BA.1 S2 (6 pro) | TCTGCTTCCTCTGTAGCTAGCCAGAGCATCATCGCCTACACCATGAGC<br>CTGGGCGCCGAGAACAGCGTGGCCTATTCCAACAACTCTATCGCCATT<br>CCCACCAACTTTACAATTAGCGTCACAACAGAGATCCTGCCCGTGAGC<br>ATGACCAAGACCAGCGTGGACTGTACAATGTACATCTGTGGCGACAGC<br>ACTGAATGCAGCAACCTGCTGCTGCAATACGGCTCCTTTTGCACCCAA<br>CTGAAGCGGGCGCTGACCGGAATCGCCGTGGAACAGGACAAAAATACC<br>CAGGAGGTGTTCGCCCAAGTGAAGCAGATCTACAAGACCCCACCTATC<br>AAGTACTTCGGCGGCTTTAACTTTAGCCAGATTCTCCCTGATCCTTCT<br>AAGCCTAGCAAGCGGAGCCCTATCGAGGATCTGCTGTTCAACAAGGTC<br>ACCCTGGCCGATGCCGGCTTTATCAAACAGTATGGCGATTGCCTGGGC<br>GACATAGCCGCCAGAGATCTGATCTGCGCCCAGAAATTCAAGGGCCTG<br>ACAGTTCTCCCACCTCTGCTGACCGACGAGATGATCGCTCAGTACACC<br>TCTGCCCTGCTGGCTGGCACCATCACATCTGGGTGGACATTTGGCGCC<br>GGCCCCGCCCTGCAGATCCCCTTTCCCATGCAGATGGCCTATAGATTC<br>AACGGAATCGGCGTGACCCAGAACGTGCTGTATGAAAACCAGAAGCTG<br>ATCGCTAACCAGTTCAATTCTGCCATCGGCAAGATCCAGGACTCCCTC<br>TCCTCTACCCCCAGCGCCCTGGGCAAACTGCAGGACGTGGTGAATCAC<br>AACGCCCAAGCCCTGAACACCCTGGTGAAGCAGCTCAGCAGCAAGTTT<br>GGCGCCATCAGCTCTGTGCTGAACGATATCTTCTCTAGACTGGACCCT<br>CCAGAAGCCGAAGTCCAGATCGATAGACTGATCACAGGCAGACTGCAG<br>TCCCTGCAAACCTACGTGACCCAACAGCTGATCAGGGCCGCTGAAATA<br>AGAGCCAGCGCCAATCTCGCCGCTACCAAGATGTCCGAGTGTGTGCTG<br>GGACAGTCTAAACGCGTTGACTTCTGCGGCAAAGGCTATCACCTGATG<br>AGCTTCCCCCAGAGCGCGCCGCACGGCGTGGTGTTCCTGCATGTGACA<br>TACGTGCCTGCCCAAGAGAAGAATTTCACAACCGCCCCTGCCATCTGC<br>CACGACGGCAAGGCCCACTTCCCAAGAGAGGGCGTTTTCGTTTCCAAT<br>GGCACACACTGGTTCGTGACACAAAGAAACTTCTACGAACCCCAGATT<br>ATCACCACCGACAACACCTTCGTGAGTGGCAATTGTGACGTGGTCATC<br>GGAATCGTGAACAACACAGTGTACGACCCTCTGCAACCTGAGCTGGAC<br>TCTTTTAAGGAAGAGCTGGACAAGTACTTTAAAAACCACACCAGCCCC<br>GATGTGGACCTGGGCGACATCAGTGGCATTAACGCCAGCGTGGTGAAC<br>ATCCAAAAGGAAATCGACAGACTGAACGAGGTGGCCAAGAACCTGAAC<br>GAGTCCCTGATCGACCTGCAGGAGCTCGGCAAATACGAGCAGGGATCC<br>GGATACATCCCCGAGGCCCCCAGAGATGGCCAGGCCTACGTGCGGAAG<br>GACGGCGAGTGGGTACTGCTGAGCACATTCCTGGGCAGATCC | SASSVASQSIIAYTMSLGAENSVAYSNNS<br>IAIPTNFTISVTTEILPVSMTKTSVDCTM<br>YICGDSTECSNLLLQYGSFCTQLKRALTG<br>IAVEQDKNTQEVFAQVKQIYKTPPIKYFG<br>GFNFSQILPDPSKPSKRSPIEDLLFNKVT<br>LADAGFIKQYGDCLGDIAARDLICAQKFK<br>GLTVLPPLLTDEMIAQYTSALLAGTITSG<br>WTFGAGPALQIPFPMQMAYRFNGIGVTQN<br>VLYENQKLIANQFNSAIGKIQDSLSSTPS<br>ALGKLQDVVNHNAQALNTLVKQLSSKFGA<br>ISSVLNDIFSRLDPPEAEVQIDRLITGRL<br>QSLQTYVTQQLIRAAEIRASANLAATKMS<br>ECVLGQSKRVDFCGKGYHLMSFPQSAPHG<br>VVFLHVTYVPAQEKNFTTAPAICHDGKAH<br>FPREGVFVSNGTHWFVTQRNFYEPQIITT<br>DNTFVSGNCDVVIGIVNNTVYDPLQPELD<br>SFKEELDKYFKNHTSPDVDLGDISGINAS<br>VVNIQKEIDRLNEVAKNLNESLIDLQELG<br>KYEQGSGYIPEAPRDGQAYVRKDGEWVLL<br>STFLGRS |

| | | |
|---|---|---|
| MERS-CoV S (2 pro) | ATGATCCATTCCGTGTTCCTGCTGATGTTTCTGTTGACTCCAACAGAG<br>AGCTACGTGGACGTAGGCCCCGACTCAGTGAAGAGCGCTTGCATCGAG<br>GTGGACATACAGCAGACGTTCTTTGATAAGACATGGCCGAGACCAATT<br>GACGTGTCTAAGGCTGACGGTATAATCTACCCACAGGGTAGAACGTAT<br>TCTAATATTACAATAACGTATCAGGGTCTCTTCCCATATCAGGGAGAT<br>CACGGGGATATGTATGTATACAGCGCTGGCCACGCCACCGGAACGACG<br>CCTCAGAAGCTCTTTGTGGCAAACTACTCCCAGGATGTAAAGCAATTT<br>GCGAATGGGTTCGTCGTTAGGATTGGAGCTGCCGCAAATTCAACGGGG<br>ACTGTTATCATTAGTCCCAGCACAAGTGCTACGATCCGCAAGATCTAT<br>CCCGCATTCATGTTGGGATCTTCAGTAGGTAACTTCAGTGATGGTAAA<br>ATGGGGAGATTCTTCAATCATACGCTTGTCCTCCTGCCGGACGGTTGC<br>GGCACCTTGCTTCGGGCATTCTACTGCATCCTCGAGCCCCGCTCCGGC<br>AACCACTGTCCCGCCGGTAATTCTTATACATCTTTCGCCACCTATCAT<br>ACGCCGGCCACTGACTGTAGCGATGGGAATTACAACAGAAATGCGAGC<br>CTGAACTCATTTAAGGAATATTTCAATCTGAGAAACTGCACGTTCATG<br>TACACTTACAACATTACTGAAGATGAGATCTTGGAGTGGTTTGGAATC<br>ACTCAGACCGCGCAGGGAGTCCACCTGTTTAGCAGCCGCTACGTTGAT<br>CTTTACGGTGGAAACATGTTTCAGTTTGCCACACTCCCCGTTTATGAT<br>ACGATAAAGTATTATTCCATTATACCTCACAGTATTCGGTCAATACAA<br>TCTGATAGGAAGGCGTGGGCAGCGTTTTATGTGTACAAACTTCAACCT<br>CTCACCTTCCTTCTTGATTTCTCTGTAGATGGTTACATAAGGAGAGCG<br>ATTGATTGTGGATTCAACGACCTTTCCCAACTGCATTGCAGCTATGAG<br>AGCTTTGACGTGGAAAGCGGCGTTTATAGTGTCAGTAGTTTCGAAGCC<br>AAACCTAGTGGAAGCGTTGTGGAGCAGGCTGAGGGCGTTGAGTGCGAT<br>TTCTCTCCTCTCCTGAGTGGAACACCTCCTCAAGTCTATAACTTTAAG<br>CGGCTTGTGTTTACAAACTGCAATTATAACCTGACCAAACTTCTTTCC<br>TTGTTTTCAGTCAATGATTTCACGTGTTCTCAGATATCACCAGCCGCG<br>ATTGCTTCAAATTGCTACTCTTCACTCATACTTGACTATTTCAGTTAC<br>CCTCTCAGTATGAAATCTGACCTTAGTGTGAGCAGTGCCGGGCCGATC<br>AGTCAGTTTAACTACAAGCAGTCCTTCAGCAACCCCACGTGTCTGATA<br>TTGGCGACAGTCCCACATAACCTTACCACGATAACCAAGCCACTCAAA<br>TACTCATACATTAACAAATGCAGCAGGTTTCTGTCTGACGACCGAACC<br>GAAGTCCCTCAGCTGGTCAACGCCAACCAATACTCTCCATGCGTGTCA<br>ATTGTCCCCTCCACTGTATGGGAAGATGGAGACTACTATCGCAAACAA<br>CTTTCACCCCTTGAGGGCGGGGGGTGGCTGGTGGCATCTGGGTCCACA<br>GTCGCTATGACGGAGCAACTCCAGATGGGATTTGGGATTACAGTCCAA<br>TACGGCACTGATACGAACTCCGTTTGTCCTAAATTGGAATTTGCTAAC<br>GACACGAAAATCGCATCTCAACTCGGAAATTGTGTAGAGTATTCCCTG<br>TATGGGGTCTCTGGTAGGGGAGTGTTCCAGAACTGTACGGCAGTTGGA<br>GTAAGGCAACAACGGTTTGTATATGACGCTTATCAGAATCTCGTGGGT<br>TACTATAGTGACGACGGCAACTATTACTGTCTGAGAGCTTGCGTCTCA<br>GTCCCCGTGAGCGTGATATATGATAAAGAAACCAAAACGCACGCTACG<br>CTCTTCGGGAGCGTAGCTTGCGAACACATAAGCAGTACGATGTCCCAG<br>TATAGTCGCTCTACCAGATCAATGCTCAAACGGCGAGATTCTACGTAC<br>GGCCCTCTTCAGACACCTGTAGGTTGCGTGCTGGGCTTGGTTAACTCA<br>AGCCTTTTTGTAGAAGATTGTAAGTTGCCTCTTGGTCAGTCCCTTTGC<br>GCCCTGCCGGACACCCCTAGCACACTTACGCCTGCCAGTGTCGGCTCA<br>GTACCAGGGGAGATGCGACTCGCTAGTATTGCTTTCAATCACCCGATA<br>CAGGTTGACCAGTTGAATTCAAGTTATTTCAAACTTTCAATTCCTACC<br>AACTTCAGTTTCGGGGTAACGCAGGAATACATTCAAACGACGATACAA<br>AAAGTCACTGTCGACTGCAAACAGTACGTATGCAACGGCTTTCAAAAG<br>TGCGAGCAATTGCTGCGGGAGTATGGTCAGTTTTGCAGTAAAATAAAT<br>CAAGCCCTTCACGGTGCGAATCTTAGACAGGACGATTCTGTTAGAAAC<br>CTTTTCGCAAGTGTCAAGTCATCTCAATCCTCACCTATTATACCAGGC<br>TTTGGGGGGGACTTTAACCTCACCCTGTTGGAGCCGGTGTCTATCTCC<br>ACGGGTTCTCGGTCCGCCCGGAGCGCTATTGAGGATCTGTTGTTCGAC<br>AAAGTCACGATAGCGGACCCTGGATATATGCAAGGGTATGATGATTGC<br>ATGCAGCAGGGTCCTGCGAGTGCGAGAGATTTGATCTGTGCGCAATAT<br>GTCGCCGGTTATAAAGTCCTCCCGCCTCTTATGGATGTCAATATGGAG | MIHSVFLLMFLLTPTESYVDVGPDSVKSA<br>CIEVDIQQTFFDKTWPRPIDVSKADGIIY<br>PQGRTYSNITITYQGLFPYQGDHGDMYVY<br>SAGHATGTTPQKLFVANYSQDVKQFANGF<br>VVRIGAAANSTGTVIISPSTSATIRKIYP<br>AFMLGSSVGNFSDGKMGRFFNHTLVLLPD<br>GCGTLLRAFYCILEPRSGNHCPAGNSYTS<br>FATYHTPATDCSDGNYNRNASLNSFKEYF<br>NLRNCTFMYTYNITEDEILEWFGITQTAQ<br>GVHLFSSRYVDLYGGNMFQFATLPVYDTI<br>KYYSIIPHSIRSIQSDRKAWAAFYVYKLQ<br>PLTFLLDFSVDGYIRRAIDCGFNDLSQLH<br>CSYESFDVESGVYSVSSFEAKPSGSVVEQ<br>AEGVECDFSPLLSGTPPQVYNFKRLVFTN<br>CNYNLTKLLSLFSVNDFTCSQISPAAIAS<br>NCYSSLILDYFSYPLSMKSDLSVSSAGPI<br>SQFNYKQSFSNPTCLILATVPHNLTTITK<br>PLKYSYINKCSRFLSDDRTEVPQLVNANQ<br>YSPCVSIVPSTVWEDGDYYRKQLSPLEGG<br>GWLVASGSTVAMTEQLQMGFGITVQYGTD<br>TNSVCPKLEFANDTKIASQLGNCVEYSLY<br>GVSGRGVFQNCTAVGVRQQRFVYDAYQNL<br>VGYYSDDGNYYCLRACVSVPVSVIYDKET<br>KTHATLFGSVACEHISSTMSQYSRSTRSM<br>LKRRDSTYGPLQTPVGCVLGLVNSSLFVE<br>DCKLPLGQSLCALPDTPSTLTPASVGSVP<br>GEMRLASIAFNHPIQVDQLNSSYFKLSIP<br>TNFSFGVTQEYIQTTIQKVTVDCKQYVCN<br>GFQKCEQLLREYGQFCSKINQALHGANLR<br>QDDSVRNLFASVKSSQSSPIIPGFGGDFN<br>LTLLEPVSISTGSRSARSAIEDLLFDKVT<br>IADPGYMQGYDDCMQQGPASARDLICAQY<br>VAGYKVLPPLMDVNMEAAYTSSLLGSIAG<br>VGWTAGLSSFAAIPFAQSIFYRLNGVGIT<br>QQVLSENQKLIANKFNQALGAMQTGFTTT<br>NEAFHKVQDAVNNNAQALSKLASELSNTF<br>GAISASIGDIIQRLDPPEQDAQIDRLING<br>RLTTLNAFVAQQLVRSESAALSAQLAKDK<br>VNECVKAQSKRSGFCGQGTHIVSFVVNAP<br>NGLYFMHVGYYPSNHIEVVSAYGLCDAAN<br>PTNCIAPVNGYFIKTNNTRIVDEWSYTGS<br>SFYAPEPITSLNTKYVAPQVTYQNISTNL<br>PPPLLGNSTGIDFQDELDEFFKNVSTSIP<br>NFGSLTQINTTLLDLTYEMLSLQQVVKAL<br>NESYIDLKELGNYTY |

| | | |
|---|---|---|
| | GCGGCCTACACGTCCTCTCTGCTTGGGAGCATTGCCGGAGTCGGATGG<br>ACTGCCGGACTTAGCAGTTTTGCGGCGATTCCATTCGCTCAGTCCATT<br>TTCTACAGACTTAACGGCGTGGGTATCACGCAGCAAGTTCTGTCTGAA<br>AACCAGAAGCTCATCGCTAACAAGTTCAATCAAGCGCTTGGGGCAATG<br>CAAACCGGTTTCACCACTACGAATGAAGCATTTCACAAAGTGCAGGAT<br>GCGGTTAATAACAATGCGCAGGCGCTTAGCAAATTGGCTTCCGAACTC<br>TCCAATACGTTTGGAGCTATAAGTGCATCAATTGGGGATATCATTCAG<br>CGGCTGGATCCCCCAGAACAAGATGCGCAGATTGACCGACTGATCAAT<br>GGGAGGTTGACGACATTGAATGCATTTGTAGCTCAGCAACTCGTGCGG<br>TCTGAGTCCGCTGCCTTGTCTGCGCAGCTTGCAAAGGATAAAGTAAAC<br>GAGTGTGTGAAGGCACAGAGTAAACGGTCCGGGTTTTGCGGCCAGGGA<br>ACGCATATTGTTAGTTTTGTTGTGAATGCGCCGAACGGCCTCTATTTC<br>ATGCACGTAGGCTACTATCCGAGCAACCATATAGAAGTAGTTTCAGCC<br>TATGGCCTGTGCGACGCTGCTAATCCCACGAACTGTATTGCTCCAGTG<br>AATGGATATTTCATCAAGACGAACAATACAAGAATCGTGGATGAGTGG<br>AGCTATACTGGCTCCTCTTTCTATGCACCTGAACCAATTACCTCTCTG<br>AACACTAAGTATGTAGCGCCGCAAGTGACCTACCAAAACATCTCTACA<br>AATCTCCCACCGCCGTTGCTTGGAAACTCCACAGGTATCGACTTTCAA<br>GATGAGCTGGACGAATTTTTCAAGAACGTAAGTACATCCATTCCCAAC<br>TTTGGGAGCCTGACACAAATCAATACGACGCTCCTCGACCTTACTTAT<br>GAGATGTTGTCCCTTCAGCAAGTAGTAAAGGCGCTTAATGAAAGTTAT<br>ATCGATCTGAAAGAGTTGGGAAACTATACTTAC | |
| MERS-CoV S2 (2 pro) | GCCAGTGTCGGCTCAGTACCAGGGGAGATGCGACTCGCTAGTA<br>TTGCTTTCAATCACCCGATACAGGTTGACCAGTTGAATTCAAG<br>TTATTTCAAACTTTCAATTCCTACCAACTTCAGTTTCGGGGTA<br>ACGCAGGAATACATTCAAACGACGATACAAAAAGTCACTGTCG<br>ACTGCAAACAGTACGTATGCAACGGCTTTCAAAAGTGCGAGCA<br>ATTGCTGCGGGAGTATGGTCAGTTTTGCAGTAAAATAAATCAA<br>GCCCTTCACGGTGCGAATCTTAGACAGGACGATTCTGTTAGAA<br>ACCTTTTCGCAAGTGTCAAGTCATCTCAATCCTCACCTATTAT<br>ACCAGGCTTTGGGGGGGACTTTAACCTCACCCTGTTGGAGCCG<br>GTGTCTATCTCCACGGGTTCTCGGTCCGCCCGGAGCGCTATTG<br>AGGATCTGTTGTTCGACAAAGTCACGATAGCGGACCCTGGATA<br>TATGCAAGGGTATGATGATTGCATGCAGCAGGGTCCTGCGAGT<br>GCGAGAGATTTGATCTGTGCGCAATATGTCGCCGGTTATAAAG<br>TCCTCCCGCCTCTTATGGATGTCAATATGGAGGCGGCCTACAC<br>GTCCTCTCTGCTTGGGAGCATTGCCGGAGTCGGATGGACTGCC<br>GGACTTAGCAGTTTTGCGGCGATTCCATTCGCTCAGTCCATTT<br>TCTACAGACTTAACGGCGTGGGTATCACGCAGCAAGTTCTGTC<br>TGAAAACCAGAAGCTCATCGCTAACAAGTTCAATCAAGCGCTT<br>GGGGCAATGCAAACCGGTTTCACCACTACGAATGAAGCATTTC<br>ACAAAGTGCAGGATGCGGTTAATAACAATGCGCAGGCGCTTAG<br>CAAATTGGCTTCCGAACTCTCCAATACGTTTGGAGCTATAAGT<br>GCATCAATTGGGGATATCATTCAGCGGCTGGATCCCCCAGAAC<br>AAGATGCGCAGATTGACCGACTGATCAATGGGAGGTTGACGAC<br>ATTGAATGCATTTGTAGCTCAGCAACTCGTGCGGTCTGAGTCC<br>GCTGCCTTGTCTGCGCAGCTTGCAAAGGATAAAGTAAACGAGT<br>GTGTGAAGGCACAGAGTAAACGGTCCGGGTTTTGCGGCCAGGG<br>AACGCATATTGTTAGTTTTGTTGTGAATGCGCCGAACGGCCTC<br>TATTTCATGCACGTAGGCTACTATCCGAGCAACCATATAGAAG<br>TAGTTTCAGCCTATGGCCTGTGCGACGCTGCTAATCCCACGAA<br>CTGTATTGCTCCAGTGAATGGATATTTCATCAAGACGAACAAT<br>ACAAGAATCGTGGATGAGTGGAGCTATACTGGCTCCTCTTTCT<br>ATGCACCTGAACCAATTACCTCTCTGAACACTAAGTATGTAGC<br>GCCGCAAGTGACCTACCAAAACATCTCTACAAATCTCCCACCG | ASVGSVPGEMRLASIAFNHPIQVDQL<br>NSSYFKLSIPTNFSFGVTQEYIQTTI<br>QKVTVDCKQYVCNGFQKCEQLLREYG<br>QFCSKINQALHGANLRQDDSVRNLFA<br>SVKSSQSSPIIPGFGGDFNLTLLEPV<br>SISTGSRSARSAIEDLLFDKVTIADP<br>GYMQGYDDCMQQGPASARDLICAQYV<br>AGYKVLPPLMDVNMEAAYTSSLLGSI<br>AGVGWTAGLSSFAAIPFAQSIFYRLN<br>GVGITQQVLSENQKLIANKFNQALGA<br>MQTGFTTTNEAFHKVQDAVNNNAQAL<br>SKLASELSNTFGAISASIGDIIQRLD<br>PPEQDAQIDRLINGRLTTLNAFVAQQ<br>LVRSESAALSAQLAKDKVNECVKAQS<br>KRSGFCGQGTHIVSFVVNAPNGLYFM<br>HVGYYPSNHIEVVSAYGLCDAANPTN<br>CIAPVNGYFIKTNNTRIVDEWSYTGS<br>SFYAPEPITSLNTKYVAPQVTYQNIS<br>TNLPPPLLGNSTGIDFQDELDEFFKN<br>VSTSIPNFGSLTQINTTLLDLTYEML<br>SLQQVVKALNESYIDLKELGNYTY |

| | | |
|---|---|---|
| | CCGTTGCTTGGAAACTCCACAGGTATCGACTTTCAAGATGAGC<br>TGGACGAATTTTTCAAGAACGTAAGTACATCCATTCCCAACTT<br>TGGGAGCCTGACACAAATCAATACGACGCTCCTCGACCTTACT<br>TATGAGATGTTGTCCCTTCAGCAAGTAGTAAAGGCGCTTAATG<br>AAAGTTATATCGATCTGAAAGAGTTGGGAAACTATACTTAC | |
| HuCoV-OC43 (2 pro) | ATGTTTCTCATCCTCCTCATTTCTTTGCCCACGGCTTTTGCGGTAATA<br>GGGGACTTGAAGTGCACCTCCGACAACATTAACGACAAGGATACTGGC<br>CCACCTCCTATCTCAACTGATACCGTTGATGTAACAAATGGGCTGGGT<br>ACGTACTACGTCCTGGATCGAGTCTATCTTAACACCACACTCTTCCTG<br>AACGGGTATTACCCTACTAGCGGCAGCACATATCGAAACATGGCTTTG<br>AAGGGATCCGTACTGCTGAGCCGCCTCTGGTTCAAACCTCCGTTTCTG<br>TCTGATTTTATCAATGGGATTTTCGCTAAGGTCAAAAACACGAAGGTA<br>ATTAAAGATCGAGTGATGTATAGCGAATTCCCGGCTATCACAATAGGC<br>TCTACTTTCGTCAATACCTCATATAGTGTGGTTGTACAACCTCGGACA<br>ATAAACTCTACGCAGGATGGCGACAATAAATTGCAAGGTCTGCTTGAA<br>GTCAGCGTGTGTCAATATAACATGTGTGAATACCCCCAGACCATCTGC<br>CATCCTAACCTTGGTAATCATCGGAAGGAGTTGTGGCACCTTGATACT<br>GGCGTAGTCTCATGCTTGTATAAGCGGAACTTTACATATGACGTAAAT<br>GCAGACTACCTCTATTTCCATTTTTACCAAGAGGGAGGCACGTTTTAC<br>GCCTACTTTACTGATACTGGCGTAGTTACGAAGTTTCTGTTTAATGTC<br>TACTTGGGAATGGCGTTGAGTCACTATTATGTTATGCCTCTCACATGC<br>AATAGTAAGCTCACCCTGGAGTACTGGGTGACTCCGTTGACATCTCGG<br>CAATATCTGCTCGCATTTAACCAAGATGGTATAATCTTTAATGCAGAG<br>GATTGTATGTCCGATTTCATGAGCGAAATTAAATGTAAAACCCAGTCT<br>ATCGCCCCACCAACAGGTGTATATGAACTCAATGGCTACACGGTGCAA<br>CCAATTGCCGATGTGTACCGACGGAAACCGAATCTGCCGAATTGTAAC<br>ATTGAAGCGTGGCTGAATGATAAAAGCGTCCCAAGCCCACTGAACTGG<br>GAGAGGAAGACATTCTCAAACTGTAACTTTAATATGTCCAGTCTGATG<br>TCTTTCATTCAAGCCGACTCCTTTACTTGCAATAATATAGATGCGGCT<br>AAAATATATGGAATGTGTTTCAGTTCCATAACTATCGATAAATTCGCC<br>ATCCCAAATGGCCGGAAGGTTGACCTTCAATTGGGGAACCTCGGCTAT<br>CTGCAATCTTTCAATTATCGCATCGACACGACAGCCACTAGCTGTCAG<br>CTTTACTATAACCTGCCGGCGGCTAACGTATCTGTTAGCCGCTTTAAT<br>CCGAGCACCTGGAACAAAAGGTTTGGTTTTATCGAAGACTCAGTTTTT<br>AAGCCTCGCCCAGCGGGCGTCCTGACTAATCACGACGTGGTGTATGCT<br>CAACATTGCTTCAAAGCCCCCAAGAACTTCTGCCCATGTAAGCTCAAT<br>GGATCATGTGTTGGTTCCGGCCCAGGGAAAAATAACGGGATCGGAACG<br>TGTCCAGCGGGAACAAACTATCTTACATGTGACAATCTTTGTACTCCT<br>GACCCAATAACATTCACAGGAACTTATAAATGTCCGCAGACGAAGAGC<br>TTGGTCGGTATAGGAGAGCACTGCTCAGGGCTGGCTGTAAAGAGTGAT<br>TACTGCGGGGGCAACAGTTGCACCTGCAGACCACAAGCTTTCTTGGGA<br>TGGTCCGCGGACAGCTGTCTGCAGGGAGATAAATGTAACATCTTTGCC<br>AATTTTATTCTTCACGACGTTAATAGCGGGTTGACCTGCTCCACTGAT<br>CTGCAAAAAGCGAACACAGATATAATCCTGGGGGTTTGCGTTAACTAC<br>GACTTGTACGGGATTTTGGGACAGGGAATCTTCGTCGAAGTAAACGCC<br>ACGTACTATAACTCCTGGCAAAACCTGCTTTATGATTCTAACGGGAAT<br>TTGTATGGGTTTAGGGACTACATCATTAATAGGACCTTCATGATAAGG<br>AGCTGTTATAGCGGTCGCGTATCTGCAGCGTTTCATGCCAACTCATCT<br>GAACCGGCACTGCTGTTCCGCAATATCAAGTGTAATTATGTTTTCAAT<br>AATTCCCTTACGAGGCAGTTGCAACCGATCAACTACTTTGACTCCTAC<br>CTGGGTTGCGTCGTAAATGCCTATAATAGTACCGCTATAAGCGTACAG<br>ACGTGTGATTTGACAGTTGGCTCCGGCTATTGCGTGGACTACTCCAAA<br>AACGGCGGTTCAGGCGGTGCGATCACCACGGGCTACCGGTTCACCAAT<br>TTCGAGCCATTTACAGTTAATTCTGTTAACGACAGCCTCGAGCCAGTC<br>GGGGGACTGTACGAAATCCAAATACCGTCAGAGTTTACAATAGGTAAT<br>ATGGTTGAATTCATACAAACCTCAAGCCCCAAGGTGACTATAGATTGC<br>GCCGCGTTTGTGTGCGGCGACTACGCTGCATGTAAAAGCCAGCTGGTG<br>GAGTATGGATCATTCTGTGACAACATCAACGCCATTCTGACTGAGGTG | MFLILLISLPTAFAVIGDLKCTSDNINDK<br>DTGPPPISTDTVDVTNGLGTYYVLDRVYL<br>NTTLFLNGYYPTSGSTYRNMALKGSVLLS<br>RLWFKPPFLSDFINGIFAKVKNTKVIKDR<br>VMYSEFPAITIGSTFVNTSYSVVVQPRTI<br>NSTQDGDNKLQGLLEVSVCQYNMCEYPQT<br>ICHPNLGNHRKELWHLDTGVVSCLYKRNF<br>TYDVNADYLYFHFYQEGGTFYAYFTDTGV<br>VTKFLFNVYLGMALSHYYVMPLTCNSKLT<br>LEYWVTPLTSRQYLLAFNQDGIIFNAEDC<br>MSDFMSEIKCKTQSIAPPTGVYELNGYTV<br>QPIADVYRRKPNLPNCNIEAWLNDKSVPS<br>PLNWERKTFSNCNFNMSSLMSFIQADSFT<br>CNNIDAAKIYGMCFSSITIDKFAIPNGRK<br>VDLQLGNLGYLQSFNYRIDTTATSCQLYY<br>NLPAANVSVSRFNPSTWNKRFGFIEDSVF<br>KPRPAGVLTNHDVVYAQHCFKAPKNFCPC<br>KLNGSCVGSGPGKNNGIGTCPAGTNYLTC<br>DNLCTPDPITFTGTYKCPQTKSLVGIGEH<br>CSGLAVKSDYCGGNSCTCRPQAFLGWSAD<br>SCLQGDKCNIFANFILHDVNSGLTCSTDL<br>QKANTDIILGVCVNYDLYGILGQGIFVEV<br>NATYYNSWQNLLYDSNGNLYGFRDYIINR<br>TFMIRSCYSGRVSAAFHANSSEPALLFRN<br>IKCNYVFNNSLTRQLQPINYFDSYLGCVV<br>NAYNSTAISVQTCDLTVGSGYCVDYSKNG<br>GSGGAITTGYRFTNFEPFTVNSVNDSLEP<br>VGGLYEIQIPSEFTIGNMVEFIQTSSPKV<br>TIDCAAFVCGDYAACKSQLVEYGSFCDNI<br>NAILTEVNELLDTTQLQVANSLMNGVTLS<br>TKLKDGVNFNVDDINFSPVLGCLGSECSK<br>ASSRSAIEDLLFDKVKLSDVGFVEAYNNC<br>TGGAEIRDLICVQSYKGIKVLPPLLSENQ<br>ISGYTLAATSASLFPPWTAAAGVPFYLNV<br>QYRINGLGVTMDVLSQNQKLIANAFNNAL<br>YAIQEGFDATNSALVKIQAVVNANAEALN<br>NLLQQLSNRFGAISASLQEILSRLDALEA<br>EAQIDRLINGRLTALNAYVSQQLSDSTLV<br>KFSAAQAMEKVNECVKSQSSRINFCGNGN<br>HIISLVQNAPYGLYFIHFSYVPTKYVTAR<br>VSPGLCIAGDRGIAPKSGYFVNVNNTWMY<br>TGSGYYYPEPITENNVVVMSTCAVNYTKA<br>PYVMLNTSIPNLPDFKEELDQWFKNQTSV<br>APDLSLDYINVTFLDL |

| | |
|---|---|
| | AATGAGCTCTTGGATACTACACAATTGCAAGTTGCTAATTCTTTGATG<br>AACGGTGTAACGCTCAGCACGAAACTCAAAGATGGTGTGAACTTTAAT<br>GTCGATGATATAAATTTCTCCCCAGTCCTTGGCTGCCTGGGCAGCGAA<br>TGTAGCAAAGCCAGTTCCCGCAGTGCTATAGAGGATTTGTTGTTCGAC<br>AAAGTAAAACTCTCTGATGTAGGGTTTGTGGAAGCATATAACAACTGC<br>ACTGGCGGTGCGGAAATTCGAGATCTTATATGCGTGCAGTCCTACAAA<br>GGAATCAAAGTCCTGCCGCCCCTGCTGAGTGAAAATCAAATCAGCGGT<br>TACACGCTCGCAGCAACTTCCGCTAGTCTTTTCCCACCTTGGACAGCT<br>GCCGCGGGCGTTCCGTTCTACCTTAATGTACAATATAGAATCAACGGG<br>CTTGGGGTAACAATGGACGTGCTTAGTCAGAACCAGAAGTTGATAGCT<br>AATGCCTTTAATAACGCTCTTTACGCAATTCAAGAAGGCTTCGACGCA<br>ACTAATTCAGCACTGGTAAAGATTCAAGCTGTAGTTAATGCCAACGCT<br>GAGGCTCTCAACAACTTGTTGCAGCAATTGAGTAACCGATTTGGGGCA<br>ATCTCAGCATCACTTCAGGAGATTTTGTCCCGATTGGATGCTCTCGAA<br>GCCGAGGCACAAATAGACCGGCTTATCAACGGTCGGCTGACCGCACTG<br>AATGCCTATGTGTCTCAACAACTTAGCGATTCTACCCTTGTAAAATTC<br>TCCGCTGCTCAAGCGATGGAAAAGGTGAATGAATGTGTGAAATCTCAA<br>TCTTCAAGAATCAATTTCTGCGGGAACGGGAATCACATTATTAGTCTG<br>GTCCAAAACGCGCCATATGGATTGTATTTCATACATTTTAGCTATGTG<br>CCTACGAAGTACGTAACTGCACGAGTGTCACCTGGACTCTGTATTGCG<br>GGGGACAGAGGGATTGCGCCGAAATCCGGATATTTCGTTAACGTTAAC<br>AATACATGGATGTATACTGGATCAGGTTACTATTATCCAGAACCTATC<br>ACCGAAAATAATGTGGTGGTTATGAGCACGTGTGCTGTAAACTACACT<br>AAGGCCCCTTATGTTATGCTTAATACGTCTATACCCAACCTTCCCGAC<br>TTCAAAGAGGAACTCGACCAGTGGTTTAAGAATCAGACGAGCGTAGCT<br>CCCGATCTTTCCCTCGATTACATTAATGTGACTTTTCTCGACCTC |
